# Combined computational and experimental analysis confirm donor-dependent optimization of critical processing parameters for improving mesenchymal stromal cell potency and expansion attributes

**DOI:** 10.64898/2026.07.03.735619

**Authors:** Oreoluwa Kolade, Kevin P. Robb, Julie Audet, Sowmya Viswanathan

## Abstract

Mesenchymal Stromal Cells (MSC) face several heterogeneity challenges hindering clinical and commercial success. Employing a multiple response model, interplay between donor heterogeneity, and critical processing parameters (CPPs), effects on MSC potency and cell expansion attributes were investigated through computed composite attribute scores. Twelve unique CPP combinations were tested in thirteen <u>m</u>arrow-derived MSC(<u>M</u>) and five <u>a</u>dipose-tissue MSC(<u>AT</u>) training and test datasets, respectively. Donor heterogeneity and select CPP conditions affected a curated gene panel (surrogate for MSC potency); while MSC expansion was primarily influenced by CPPs. Model performances were evaluated against clinical effectiveness data from a previously deployed clinical trial; top-performing model predicted donor rankings coincided with clinical effectiveness data, validating the modeling approach used. Our model predicted that only 8% of tested donors were agnostic to CPPs; a majority (62%) of donors showed CPP-dependent optimal composite quality attributes, with MSC seeding density as a key driver; medium supplementation and oxygen preferences were highly donor dependent. Approximately 30% of donors performed poorly at all conditions tested and may be prospectively identified using a subset of genes (***TGFB, VEGF, PDCD1LG1, PDCD1LG2, IDO***). Model predicted optimal parameters worked for 69% of tested donors, while sub-optimal parameters worked for only 23% of donors and were confirmed in an independent CD14^+^ macrophage assay. Our integrated computational and experimental framework predictably identified interactive effects of donor heterogeneity and CPP conditions to optimize MSC potency attributes.

## 1 Introduction

Mesenchymal stromal cells (MSC) with reported immunomodulatory and regenerative properties have been promising candidates for various clinical applications^1–4^. Despite this potential, the translation of MSCs into approved therapies has been challenging. Remestemcel-L, an allogenic MSC(M) was approved by the Food and Drug Administration (FDA), after some 30 years of iterative clinical development^5^. Another MSC product Darvadostrocel, an allogeneic adipose tissue MSC(AT) originally approved in 2017 to treat complex perianal fistulas in patients with Crohn’s Disease was voluntarily withdrawn after failing to show significant differences between treatment and placebo groups in larger, powered Phase III trials^6^.

Primary tissue donor heterogeneity remains a major challenge in MSC clinical translation. While early-stage trials using limited number of donors show promising results, advanced trials using a larger cohort of donors often underperform^7–11^; a key reason is batch-to-batch variance arising from use of multiple primary tissue donors^7^ given that primary MSCs senesce in long-term cultures^12^ and are used clinically at population doublings ≤ 20^14^. This necessitates multiple primary tissue donors to satisfy clinical dosing needs, exacerbating MSC heterogeneity issues. In fact, approval for Remestemcel-L MSC product was only obtained after implementing process improvements that reduced batch-to-batch heterogeneity as demonstrated by potency attributes^13^. Intrinsic donor differences including donor age, sex, epigenetics are further compounded by donor-specific sensitivity to manufacturing conditions^14–15^ and batch-to-batch variability^16–17^. Recipient host microenvironmen**t**^18–19^ and patient baseline immunophenotypic status, addressed elsewhere further contribute to heterogeneity in clinical effectiveness^20^

MSC potency attributes often lack sensitivity to donor heterogeneity and can therefore fail to predict batch-to-batch variations^7^, which contributes to clinical inefficacy. A new generation of quantifiable critical quality attributes (CQAs) that can be correlated with clinical efficacy are emerging^21–22^. However, donor heterogeneity is still addressed in silos with only a few studies examining its effects on MSC potency attributes^23^; the effects of manufacturing conditions on MSC potency and cell expansion are also studied^24–25^, but again without integrating donor heterogeneity effects.

The interactive effects of donor heterogeneity and manufacturing conditions on MSC potency and cell expansion attributes thus, remain a gap, and are addressed here using a systematic statistical modelling approach. Combining experimental data and multi-response desirability analysis^26^, the generated model predicts subcategories of donors i) associated with high potency that were validated against previous clinical effectiveness^27^; and ii) donor categories that required donor-specific CPP optimization; all model predictions were tested in an independent test dataset of adipose-tissue MSC(AT) and against an independent immumodulatory functional assay. Multi-response model predictions revealed optimal CPP conditions, both donor-dependent and-independent that can be used to mitigate MSC(M) donor-driven inconsistencies.

## 2 Materials and Methods

### 2.1 MSC(M), MSC(AT) Donors, and CPP Variables

Intrinsic donor heterogeneity was investigated using thirteen MSC(M) donors, including a mix of ten diseased, Osteoarthritis (OA) donors, bio-banked from a previously conducted clinical trial^28^ and three non-diseased donors (UHN REB 14-7843). All experiments were conducted at passages 5-7. Donor demographics for a subset of ten OA MSC(M) donors were previously reported^27^; two donors were excluded due to unavailability. Donor MSC(M) and MSC(AT) doubling time (T_d_), gene expression measurements (ΔCt), and CPP variables are described in *Supplementary Files*.

### 2.2 MSC(M) Training Dataset Donors Classifications

A subset of MSC(M) training dataset donors (10 of 13) had clinical efficacy data assessed at 12-months using the Knee Injury and Osteoarthritis Outcome Score (KOOS) for pain and function relative to baseline, reported as percent changes; Function-Pain Responders were defined based on meeting predetermined thresholds^29^ of improvements in *both* pain and function^27^.

### 2.3 Multiple Response Model

A multiple response model was used to evaluate how MSC attributes through ΔCt and T_d_ values were jointly influenced by the interplay between donor heterogeneity and CPPs. Model details are provided in *Supplementary Files*.

### 2.4. TNF soluble factor expression

TNF soluble factor production was measured in human macrophages, differentiated from CD14^+^ peripheral monocytes over 5 days with 20 ng/mL M-CSF (PeproTech). Macrophages (100,000 cells/well) from mismatched healthy donors (UHN REB 14-7483) were cultured as before^30^. For a subset of donors and CPP combinations, MSC(M) or MSC(AT) were cultured as described in the *Supplementary Files*, under specific CPP condition, without cytokine licensing; the resulting, CPP-specific conditioned medium (CM) was collected after 7 days. CM was added to macrophages at 30% (v/v%) for an additional two days; control wells received only basal medium. 2.5 ng/mL of lipopolysaccharide (LPS) was added 4 hours prior to readout. As controls, macrophages were polarized for 2 days towards pro-resolving with (IL-10, TGFβ, 20ng/mL each) or pro-inflammatory (IFNγ 100ng/mL; LPS, 2.5ng/mL) phenotypes.

### 2.5 Statistical Analysis

Data was visualized with GraphPad Prism 12 and JMP®, with a range of statistical analyses employed including Tukey Honest Significant Difference (HSD) test; Zero-Inflated Sinh-Arc Sinh (ZI SHASH) distribution, Kruskal-Wallis non-parametric test; hieratical clustering; Principal Component Analysis (PCA); cumulative probability distribution model and Spearman’s correlations. Assumptions for the HSD test included data normality and homoscedasticity. Receiver operating characteristic (ROC) analysis with area under the curve (AUC) was used to evaluate classification performance across all thirteen iDS models and discriminative ability of the CPPs against a binary TNF outcome, as detailed in the *Supplementary Files*. AUC values were estimated using DeLong 95% confidence intervals, and CPP-outcome associations were assessed via Fisher’s Exact test. Statistical significance was defined as p<0.05, with multiple comparisons assessed via Tukey’s test for parametric datasets.

## 3 Results

### MSC(M) donor heterogeneity and CPP variations impact potency and expansion CQAs

To interrogate the interplay between <u>m</u>arrow-derived MSC<u>(M</u>) intrinsic donor heterogeneity and CPP combinations on MSC cell expansion and potency, we identified three main CPP conditions, selected for ease of manipulation and known effects on MSC expansion^31;^ these included cell seeding density (CD), medium composition (MC), and oxygen (O_2_) concentrations, resulting in 12 unique CPP combinations. 13 non-diseased and diseased MSC(M) donors formed the training dataset (Suppl. Table 1). Two key MSC(M) CQAs were evaluated, including a curated panel of eight immunomodulatory and angiogenic genes (Suppl. Table 2), selected based on correlations with MSC(M) responses in knee OA patients^21^ and previous evidence^32–38^ as surrogate measures of MSC(M) potency. Multivariate gene expression profiles are relatively high throughput and correlate with immunomodulatory and angiogenic functionality^21,39^. MSC(M) cell expansion, assessed as doubling times (T_d_) over 7 days, served as the second CQA and resulted in a median T_d_ of 3.85 (95% CI: 3.69 - 4.59; Suppl. Table 3). A 7-day timeframe allowed for relatively high throughput readouts to test multiple CPP and MSC donor combinations.

Unsupervised hierarchical clustering of gene expression for 13 MSC(M) donors under 12 CPP combination demonstrated separation based on MSC(M) donor heterogeneity (Fig. 1A). PCA of a subset of donors (10 of 13) tested under one CPP condition, used previously in a clinical trial^28^ showed segregation of donors based on clinical effectiveness into Function-Pain Responders vs. Non-Responders (Fig S1A-C), as described^27^. PC1 drove 62.5% of the variance; immunomodulatory genes (*TGFb, VEFG*, and *PDCD1LG2*) primarily contributed to PC1 variance (Fig S1D); although angiogenic genes *H1F-1a* and *TWIST1* also contributed to PC1, their relative expression was significantly lower (Fig S1E). Importantly, there was significantly higher immunomodulatory gene expression in MSC(M) donors classified as Function-Pain Responders vs. Non-Responders; *TNFA1P6*, although not significant, trended higher in Function-Pain Responders (Fig S1E). AUC using the mean of direction-corrected gene expression for five genes (*TGFB, VEGF, PDCD1LG1, PDCD1LG2, IDO*; Fig S1F) associated with PC1, was able to discriminate all 10 donors based on previous clinical performance with 100% accuracy^27^.

**Figure 1:**
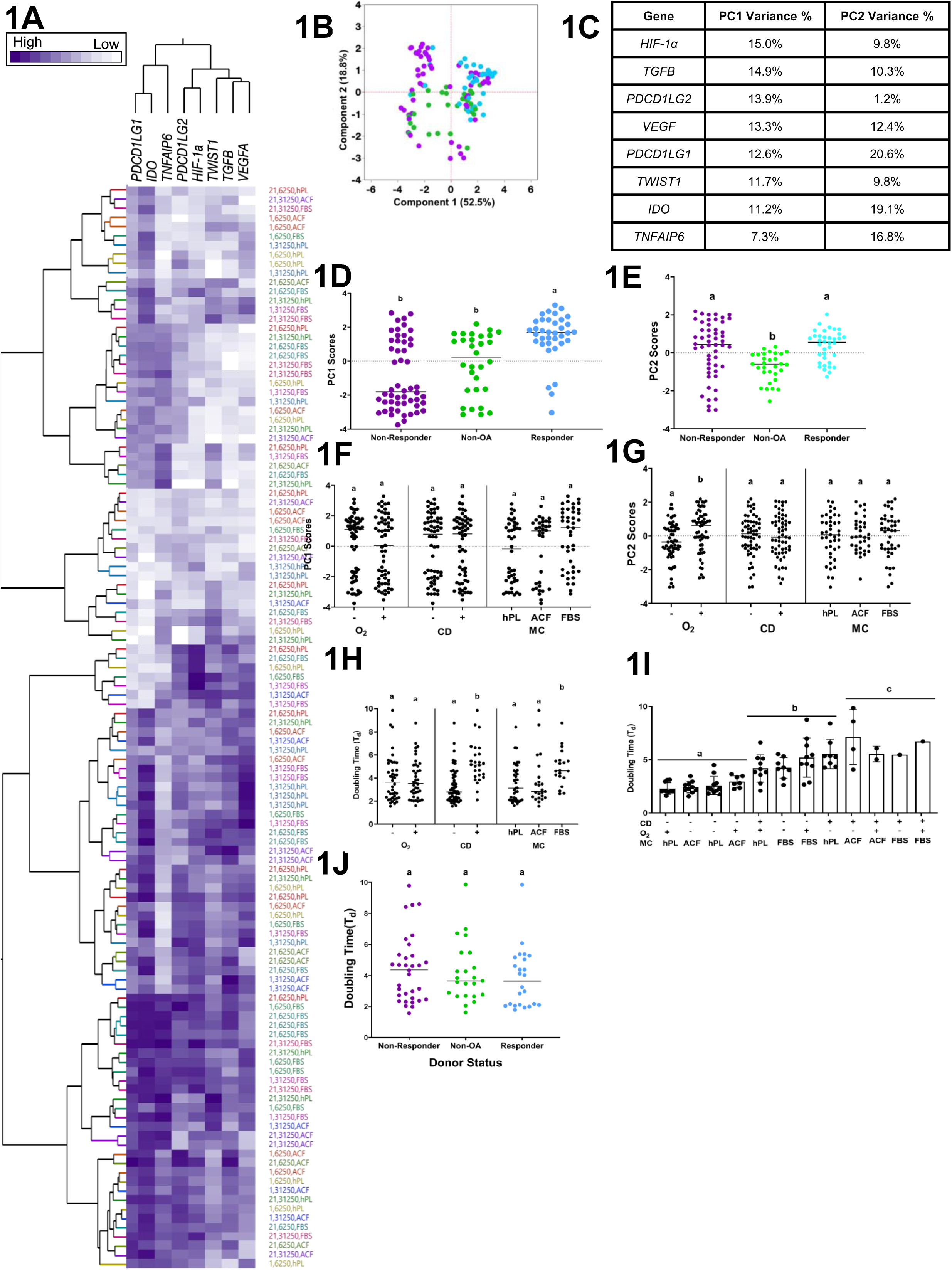
Curated gene panel is a sensitive discriminator of donor heterogeneity while doubling time (T_d_) is sensitive to variations in CPPs. (**A**) Unbiased hierarchical clustering (Ward Method) of all CPP conditions (colour-coded dendrogram) and all MSC(M) training set donors (n=13). (**B**) Score plot using PCA clustering shows separation by donor categorization with Function-Pain Responders^27^ shown in blue, Non-Responders in purple, and Non-OA Donors in green. (**C**) Contribution of eight curated genes to PC1 and PC2 % variance. (**D**) PC1 and (**E**) PC2 scores across donor groups. (**F**) PC1 and (**G**) PC2 scores across CPP conditions; specifically O_2_ at 1% (–) or 21% (+); CD at 6,250 cells/cm² (–) or 31,250 cells/cm² (+); MC including ACF, hPL, or FBS. (**H**) T_d_ across CPP categories. (**I**) T_d_ measurements across 13 donors and indicated CPP combinations (**J**) T_d_ across donor categories. Lowercase letters indicate significant pairwise differences based on Tukey’s HSD test. Error bars represent mean±SD (Donor n =13). Group differences across CPP conditions and donor categories were evaluated using ANOVA followed by Tukey’s HSD post-hoc test. *ACF- Animal component free; FBS- Fetal bovine serum; HIF-1a-Hypoxia-inducible factor 1-alpha; hPL- Human platelet lysate; HSD- Honestly Significant Difference; IDO- Indoleamine 2,3-dioxygenase; PC1- Principal component 1; PCA- Principal Component Analysis; PDL1- Programmed death-ligand 1; SD- Standard Deviation; TGF* β*-transforming growth factor beta; TNFAIP6- tumour necrosis factor-alpha stimulated gene; TWIST1- Twist-related protein1; VEGF- Vascular endothelial growth factor.*

This donor categorization thus was extrapolated to all training set donors (n=13), across 12 CPP conditions. PC1 scores (Fig. 1B) now captured 52.5% of total variance, and was still dominantly driven by expression of *TGFb, PDCD1LG2, VEGF* (Fig. 1C). PC1 scores remained significantly higher in Function-Pain Responders vs. Non-Responders (p<0.05); non-diseased donors, now added, exhibited intermediate PC1 scores, with statistically lower PC1 scores vs. Function-Pain Responders (p<0.05; Fig. 1D). Donor heterogeneity was also reflected in PC2 Scores with both Function-Pain Responders and Non-Responders exhibiting significantly higher gene expression than non-diseased donors (Fig. 1E).

The curated genes in PC2, but not PC1 were sensitive to variations in CPP conditions (Fig. 1F, Fig S1G-I), specifically to CPP conditions associated with O_2_ concentration (Fig 1G). T_d_ also showed sensitivity to CPP conditions, particularly CD and MC levels (p<0.05; Fig. 1H); shorter T_d_ was significantly associated with low CD, typically with human platelet lysate (hPL) or animal component free (ACF) medium supplementation, regardless of O_2_ levels. Conversely, longer T_d_ occurred at high CD, in ACF- and FBS- supplemented conditions, again, agnostic to O_2_ concentrations (Fig. 1I); donor category had no significant impact on T_d_ (Fig. 1J). These findings underscore the advantage of using xenofree medium supplementation and lower CD to enhance MSC(M) expansion. This finding was generalizable to other MSCs as confirmed in MSC (AT) test dataset (Fig. S2).

### A framework to study interaction effects between MSC donor heterogeneity and CPP conditions

While the PCA was useful in confirming that the chosen MSC potency and cell expansion readouts were sensitive to CPP and donor heterogeneity, it does not allow for assessment of their interaction effects. To address this, a linear multi-response model that incorporate interaction effects of donor heterogeneity and CPP on MSC potency and cell expansion was developed. The multi-response model using 13 MSC(M) donors x 12 CPP conditions identified significant main effects for CD, donor, MC, O□, and batch effects (Fig 2A); CD and donor variation exerted the greatest influence on the multi-response model; significant interaction effects were also identified between donors x O_2_ and O x MC (Fig. 2A).

**Figure 2:**
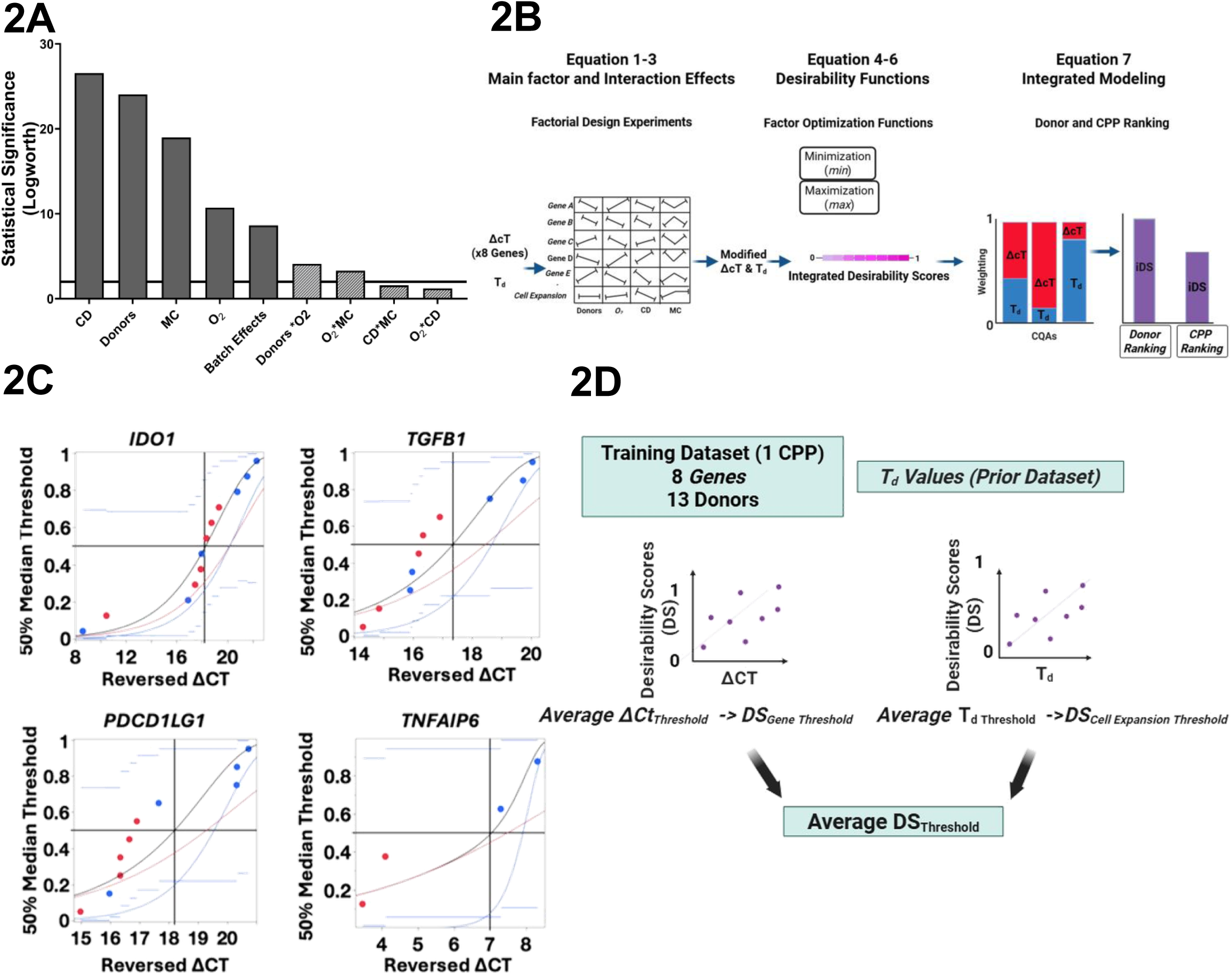
Multi-response modeling workflow and determination of iDS. (**A**) Effects summary table showing the statistical significance of main effects and donor interaction terms; solid horizontal line denotes Logworth corresponding to p<0.05. Logworth = −log□□ (p). Dark bars represent main effects (CD, Donors, MC, O□, Batch Effects); lighter bars represent two-factor interaction effects (Donors×O□, O□×MC, CD×MC, O□×CD). **(B)** Schematic overview of the multi-response model workflow for iDS generation (**C**) ΔCT values from 10/13 MSC(M) training set donors associated with a clinical response^28^ for four genes (*IDO1, PDCD1LG1, TGFB1, TNFAIP6*) were classified using a 50% median. Horizontal blue lines represent 95% confidence intervals for mean or median parameters. The y-axis denotes culmulative probability, and the x-axis shows reversed ΔCT values (empirically adjusted as (–1(ΔCT) + 20), with increasing expression from left to right. (**D**) Schematic of the iDS threshold determination workflow. ΔCT_Threshold_ values from full MSC(M) training dataset donors under one CPP condition (6250 cells/cm², 21% O□, FBS) were converted to DS_Gene Threshold_ via regression equations (*Suppl. Table 9*). T_d_ values from a previously published reference dataset^28^ were independently used to identify DS_Cell Expansion Threshold_ via regression equations (*Suppl. Table 10*). DS values from DS_Gene Threshold_ and DS_Cell Expansion Threshold_ were then averaged to define the final threshold of DS_Threshold_. - Δ*CT- negative delta cycle threshold; iDS- Integrated Desirability Scores; DS-Desirability Scores*.

To explore the main and interactive effects of donor heterogeneity and CPPs on MSC(M) gene expression and cell expansion, a four factor multi-response model was developed using “Donor” (13 donors); O_2_ (2 levels: normoxic, hypoxic); CD (2 levels: 6250 cells/cm^2^, 31250 cells/cm^2^) and MC (3 levels: ACF, FBS, hPL; Suppl. Table 4). Multi-response optimization functions (Equation 1-3; *Supplementary Files*) calculated interaction coefficients between these four-input parameters (Fig 2B). The interaction coefficients were applied to experimentally measured ΔCt and T_d_ values to generate adjusted ΔCt or T_d_ values by adding model-estimated main effect and interactions terms (Suppl. Table 5). The adjusted ΔCt and T_d_ values were transformed via regression into unitless desirability scores (DS) for both gene expression (DS_genes)_ and cell expansion (DS_cell expansion_) using minimization and maximization functions (Equation 4–6, *Supplementary Files;* Fig 2B). These DS_gene_ and DS_cell expansions_ were integrated into composite attribute scores, called the integrated desirability score (iDS, from 0 to 1) using different model weightings (Equation 7, *Supplementary Files;* Fig 2B). Empirical model weightings allowed assessments of the relative contribution of gene expression vs. cell expansion to overall MSC composite quality index, calculated as iDS (Suppl. Table 6,7). iDS values were generated across the full parametric space, including unsampled conditions, enabling model-predicted ranking of donor heterogeneity and identification of optimal CPP combinations. Model-predicted iDS were compared to independent test datasets (Fig 2B).

To interpret the model outputted iDS, we had to define what was “desirable.” ΔCt values for four genes shared between the current MSC(M) training and a previous MSC(M) dataset with clinical effectiveness data^28^ (*IDO, PDCD1LG1, TGFB1, TNFAIP6)*, were modelled using a cumulative probability distribution, fitted to a smallest extreme value (SEV) distribution (Fig 2C). As 10 of 13 donors from the training dataset overlapped with the previous MSC(M) dataset, Function-Pain Responder vs. Non-Responder donor classifications for these 10 donors^27^ were used as failure analogs and correlated with changes in previously measured ΔCt values. A 50% median resulted in low misclassification rates for three of the four genes (0%, 22.2%, 22.2% for *TNFAIP6*, *PDCD1LG1*, *TGFβ* respectively); *IDO1* however, showed poorer performance with only 50% correct classification (Suppl. Table 8).

Given that the 50% median had a relatively low misclassification rate for three of the four genes, this median was applied to all eight genes (Fig 2D). Raw, non-modified, experimentally-derived ΔCt values (without consideration for interaction effects to establish an unbiased threshold) from the full MSC(M) training dataset, associated with the 50% median, were transformed into DS_genes_ values. The mean of eight DS_genes threshold_ (0.556) was used as a threshold for desirable gene expression (Suppl. Table 9).

Similarly, non-modified T_d_ values from the previous dataset (only 10 of 13 MSC(M) training dataset donors included; Suppl. Table 10) were converted to equivalent DS_cell expansion threshold_ values; an average DS_cell expansion threshold_ of 0.427 days was obtained; doubling times longer than this were considered undesirable. An integrated threshold, iDS_threshold_ was generated by summing the DS_gene threshold_ and DS_cell expansion threshold_. These thresholds could be combined using different weightings of the gene expression: cell expansion, specifically, 100:0%; 90:10%; 80:20%; 50:50% and 0:100% weightings were empirically assessed. These produced iDS_thresholds_ of 0.556, 0.543; 0.530; 0.492, and 0.427 respectively. For subsequent analyses, the 80:20 weighting scheme, corresponding to an iDS_threshold_ of 0.530 ± 0.086, was empirically selected. With this weighting scheme, 39% of donor-CPP combinations exceed the desirability criteria (Suppl Table 11); toggling the weighting schemes to 100:0 or 0:100 resulted in 24% or 79% of donor-CPP combinations meeting desirability criteria (Suppl Table 11). Desirable iDS were defined as those exceeding 0.530; non-desirable iDS fell below 0.444 (0.530 - SD); iDS within this range (0.444-0.530) were classified as rescuable.

### Multi-response model performance verified against clinical effectiveness

The multi-response model tested different weightings for MSC(M) gene expression vs. cell expansion potency attributes. 13 predictive models of iDS were generated using different weightings to rank MSC(M) training set donors (Table 1; Suppl. Table 11) and to identify optimal CPP combinations (Table 2). Models 1, 2, 3, 4 and 5 corresponded to DS_gene_:DS_cell expansion_ weightings of 90:10, 80:20, 50:50, 100:0 and 0:100 respectively; models A, B, and C represented sub-weightings based on eight genes; three most sensitive genes; and eight genes with orthogonal directionality assigned to immunomodulatory vs. angiogenic genes respectively (Suppl. Table 7).

**Table 1:** Model predicted donor iDS rankings vs. donor rankings, based on previous clinical performance. Model predicted iDS-based donor rankings for a subset of MSC(M) training dataset donors (10/13) across 13 models, generated by evaluating different ΔCT:T_d_ % weighting schemes (Model 1: 90:10; Model 2: 80:20; Model 3: 50:50; Model 4: 100:0; Model 5: 0:100). Within each ΔCT inclusive scheme (Models 1,2,3 and 4), further sub-weightings were performed for the 8 curated genes as Model A (all genes); Model B (top three most sensitive genes); Model C (directionally weighted, penalizing conditions increasing *HIF1A, TWIST1*, or *VEGFA* and rewarding upregulation of other genes). Actual donor rankings based on combined percent-change KOOS Function and Pain scores from baseline to 12 months, as reported^27^. AUC was calculated to quantify the ability of each model to classify Function-Pain Responders and Non-Responders using the continuous IDS scores generated by the model. Spearman correlations between model-predicted iDS vs. KOOS outcomes, with significant p-values, corresponding AUC and rho correlation values are bolded. *ADL-Activity of Daily Living (Function score); CPP- Critical Process Parameter; iDS- Integrated Desirability Scores; KOOS- Knee injury and Osteoarthritis Outcome Score; T_d_ - Doubling Time*

|  | Actual Ranking of<br>OA Donor ID | Model predicted donor rankings based on iDS |  |  |  |  |  |  |  |  |  |  |  |  |
| --- | --- | --- | --- | --- | --- | --- | --- | --- | --- | --- | --- | --- | --- | --- |
|  |  | Model<br>1A | Model<br>1B | Model<br>1C | Model<br>2A | Model<br>2B | Model<br>2C | Model<br>3A | Model<br>3B | Model<br>3C | Model<br>4A | Model<br>4B | Model<br>4C | Model<br>5 |
|  | 7 (100) | 7 | 7 | 4 | 7 | 7 | 4 | 7 | 7 | 4 | 7 | 7 | 4 | 1 |
|  | 11 (75.69) | 5 | 6 | 6 | 5 | 6 | 6 | 5 | 5 | 5 | 6 | 6 | 6 | 5 |
|  | 13 (70.38) | 6 | 5 | 5 | 6 | 5 | 5 | 11 | 6 | 6 | 5 | 5 | 5 | 4 |
|  | 5 (64.72) | 11 | 11 | 7 | 11 | 11 | 7 | 6 | 11 | 7 | 13 | 11 | 7 | 11 |
|  | 10 (31.00) | 13 | 13 | 10 | 13 | 13 | 10 | 2 | 2 | 10 | 11 | 13 | 10 | 7 |
|  | 1 (30.65) | 2 | 2 | 11 | 2 | 2 | 11 | 13 | 13 | 11 | 2 | 2 | 11 | 10 |
|  | 2 (-3.47) | 10 | 10 | 1 | 10 | 10 | 1 | 10 | 10 | 1 | 10 | 10 | 1 | 6 |
|  | 4 (-24.11) | 4 | 4 | 2 | 4 | 4 | 2 | 4 | 4 | 2 | 4 | 4 | 2 | 12 |
|  | 12 (-61.89) | 1 | 12 | 12 | 1 | 1 | 12 | 1 | 1 | 12 | 1 | 12 | 12 | 2 |
|  | 6 (-83.30) | 12 | 1 | 13 | 12 | 12 | 13 | 12 | 12 | 13 | 12 | 1 | 13 | 13 |
| Model<br>Performance | AUC | **0.89** | 0.87 | 0.52 | **0.85** | 0.82 | 0.52 | 0.66 | 0.64 | 0.50 | **0.90** | 0.84 | 0.52 | 0.50 |
|  | rho correlation | **0.72** | 0.60 | -0.08 | **0.72** | 0.55 | 0.03 | 0.50 | 0.47 | 0.21 | **0.74** | 0.59 | 0.11 | 0.30 |
|  | p-value | **0.01** | 0.07 | 0.83 | **0.02** | 0.10 | 0.93 | 0.14 | 0.17 | 0.56 | **0.01** | 0.07 | 0.75 | 0.40 |

**Table 2:** Donor-specific MSC(M) iDS across CPP conditions. CPP combinations were evaluated across 13 MSC(M) training dataset donors using the top-performing model weightings (Models 1A). Donor-specific iDS values were reported for each CPP condition, with corresponding mean iDS calculated across all tested CPPs. CPPs with iDS ≥ 0.530 classified as desirable and bolded.

| CPPs | OA Donor 7 | OA Donor 5 | OA Donor 6 | OA Donor 11 | OA Donor 13 | Non-OA Donor 2 | OA Donor 2 | Non-OA Donor 1 | Non-OA Donor 3 | OA Donor 10 | OA Donor 4 | OA Donor 1 | OA Donor 12 | Mean iDS per CPP | No. Donors that exceed Threshold (No. Donors that do not) |
| --- | --- | --- | --- | --- | --- | --- | --- | --- | --- | --- | --- | --- | --- | --- | --- |
| 1,6250,FBS | **0.70** | **0.62** | **0.73** | **0.63** | **0.67** | **0.60** | **0.58** | **0.56** | **0.63** | 0.51 | 0.49 | 0.43 | 0.40 | **0.58**<sub>0.10</sub> | 9(4) |
| 21,6250,hPL | **0.72** | **0.69** | **0.57** | **0.64** | **0.61** | **0.58** | **0.56** | **0.56** | 0.37 | 0.43 | 0.39 | 0.36 | 0.32 | 0.52<sub>0.13</sub> | 8(5) |
| 21,6250,FBS | **0.70** | **0.67** | **0.55** | **0.62** | **0.58** | **0.58** | **0.55** | **0.56** | 0.43 | 0.43 | 0.40 | 0.36 | 0.34 | 0.52<sub>0.12</sub> | 8(5) |
| 21,31250,hPL | **0.71** | **0.68** | **0.56** | **0.63** | **0.60** | **0.60** | **0.56** | **0.56** | 0.40 | 0.39 | 0.39 | 0.36 | 0.33 | 0.52<sub>0.13</sub> | 8(5) |
| 1,6250,ACF | **0.64** | **0.54** | **0.66** | **0.56** | **0.60** | **0.54** | 0.51 | 0.48 | **0.54** | 0.42 | 0.39 | 0.35 | 0.31 | 0.50<sub>0.11</sub> | 7(6) |
| 1,31250,FBS | **0.60** | **0.54** | **0.63** | **0.55** | **0.58** | 0.48 | 0.50 | 0.50 | **0.55** | 0.43 | 0.42 | 0.38 | 0.34 | 0.50<sub>0.09</sub> | 6(7) |
| 1,6250,hPL | **0.61** | 0.51 | **0.63** | **0.53** | **0.56** | 0.50 | 0.47 | 0.44 | 0.52 | 0.40 | 0.36 | 0.31 | 0.27 | 0.47<sub>0.11</sub> | 3(10) |
| 1,31250,ACF | **0.58** | 0.51 | **0.60** | 0.51 | **0.54** | 0.47 | 0.46 | 0.46 | 0.51 | 0.40 | 0.39 | 0.34 | 0.29 | 0.47<sub>0.09</sub> | 3(10) |
| 21,6250,ACF | **0.66** | **0.62** | 0.50 | **0.58** | **0.56** | **0.53** | 0.51 | 0.52 | 0.36 | 0.36 | 0.28 | 0.28 | 0.26 | 0.46<sub>0.14</sub> | 5(8) |
| 21,31250,ACF | **0.63** | **0.59** | 0.48 | **0.56** | 0.48 | 0.52 | 0.49 | 0.50 | 0.37 | 0.36 | 0.33 | 0.30 | 0.28 | 0.45<sub>0.11</sub> | 3(10) |
| 1,31250,hPL | **0.58** | 0.48 | **0.60** | 0.50 | **0.53** | 0.47 | 0.44 | 0.42 | 0.49 | 0.38 | 0.35 | 0.30 | 0.26 | 0.45<sub>0.10</sub> | 3(10) |
| 21,31250,FBS | **0.62** | **0.58** | 0.47 | **0.55** | 0.42 | 0.52 | 0.47 | 0.49 | 0.39 | 0.35 | 0.34 | 0.29 | 0.28 | 0.44<sub>0.11</sub> | 3(10) |

For a subset of donors (10 of 13), model predictions of donor rankings, based on iDS were compared to donor performance from a prior clinical trial^28^(Table 1). Discriminative performance of each model was assessed by ROC analysis, using continuous iDS as the predictor variable and Function-Pain Responder/Non-Responder status as binary outcomes (Fig 3A). Top-performing models included 4A, 1A and 2A, which had significant AUCs of 0.90 (95% CI 0.85-0.96), 0.89 (0.83-0.95), 0.85 (0.78-0.92) respectively (Suppl. Table 12). These models also exhibited significant Spearman’s correlations between iDS and percent improvements in both 12-month KOOS function and pain scores (Table 1); correlations were not adjusted for covariates, as previously justified^27^. All three models were able to significantly discriminate between different categories of donors (Fig 3B, C, D). From these, Model 1A was selected for subsequent analysis, as it combined gene expression and cell expansion metrics (Model 4A only considered gene expression), and performed slightly better than Model 2A. Leave-out-one (LOO) sensitivity analysis with Model 1A indicated that correlations were robust and not significantly skewed by omitting any single donor (Suppl. Table 13).

**Figure 3:**
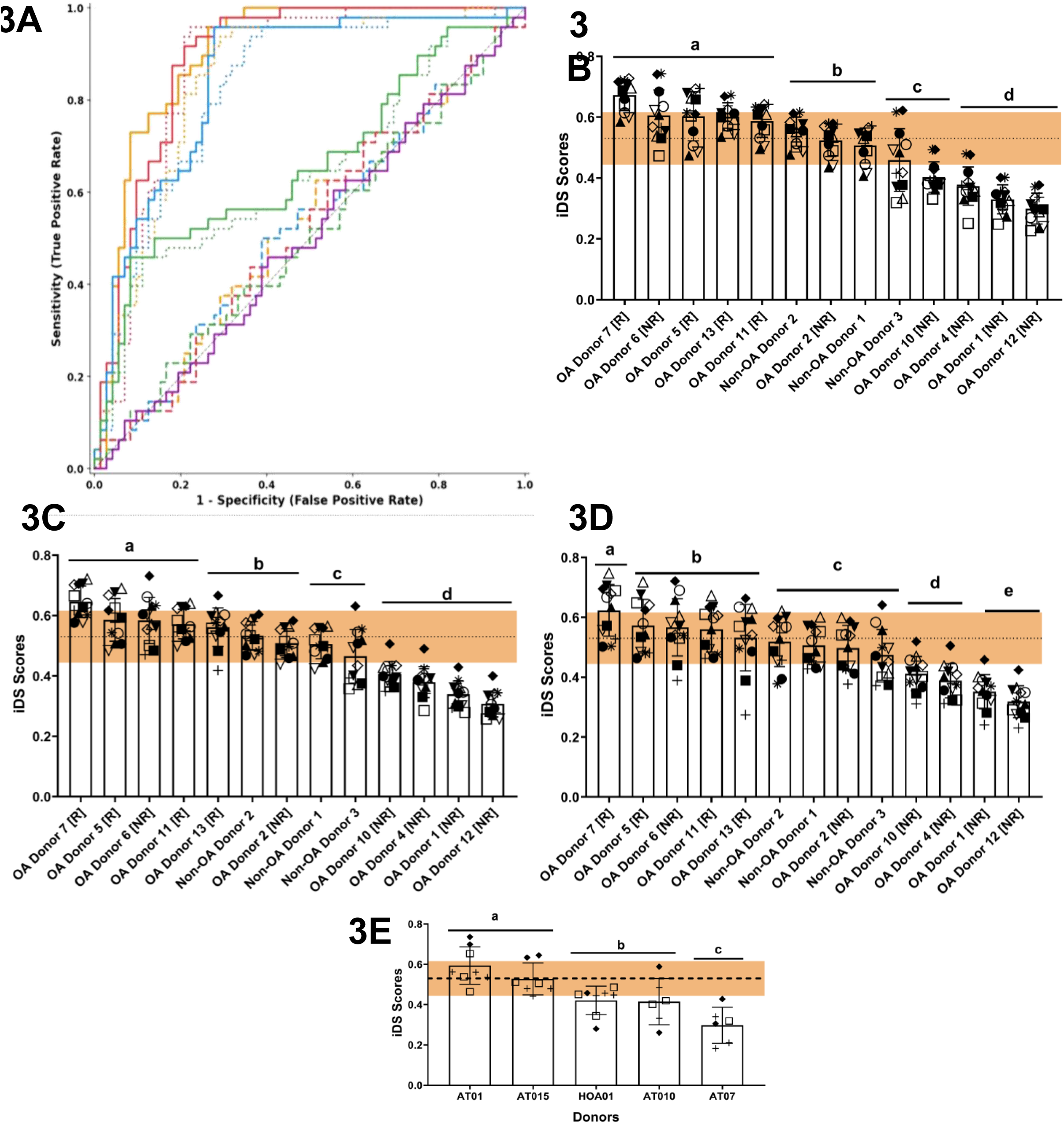
Model-predicted iDS across donors for MSC(M) training and MSC(AT) test datasets. **(A)** ROC curves for all 13 models across the full MSC(M) training dataset. Models 1,2,3,4,5 are distinguished by colour (red, blue, green, orange, and purple, respectively), with line style denoting sub-models (solid = A, dotted = B, dashed = C). The diagonal dashed line represents chance-level discrimination**. (B–D**) Model-predicted iDS for MSC(M) training dataset using Models 4A, 1A and 2A respectively. (**E**) Model-predicted iDS using Model 1A in MSC(AT) test dataset. The experimentally determined iDS threshold using an 80%:20% ΔCT:DT weighting scheme (dotted line) was 0.530, with a standard deviation of ±0.086 (range shown in orange). MSC(M) training dataset with donors categorized into OA donors (1 -13) that were further sub-categorized as responders (R), non-responders (NR) based on previous classification of clinical responsiveness^27^, and Non-OA donors (1–3). 12 CPP combinations including: (O) for low CD/ACF/low O_2_, (□) for low CD/ACF/high O_2_, (●) for high CD/ACF/low O_2_, (◼) for high CD/ACF/high O_2_, (▲) for low CD/hPL/low O_2_, (□) for low CD/hPL/high O_2_, (□) for high CD/hPL/low O_2_, (▼) for high CD/hPL/high O_2_, (□) for low CD/FBS/low O_2_, (□) for low CD/FBS/high O_2_, (*) for high CD/FBS/low O_2_, and (+) for high CD/FBS/high O_2_. Lower case letters indicate significant differences between specific groups of donors using Tukey’s HSD test.

Model 1A performance was also tested for its ability to rank donors in an independent test dataset of five MSC(AT) donors under three CPP conditions (1%/6250/FBS, 21%/6250/ACF and 21%/31250/FBS, Fig 3E). Predicted iDS rankings corresponded with previously reported donor potency rankings^21^ based on immunomodulatory functionality (Suppl. Table 14). Thus, the multi-response model accurately identified potent MSC donors across both training and independent test datasets.

### Model predictions of optimal CPP combinations show donor dependency

Next, we evaluated the multi-response model predictions for optimal CPP combinations focusing only on the top performing Model 1A. For each MSC(M) training set donor, model predicted iDS ranked 12 CPP combinations (Table 2). The top-performing CPP combination across all donors was 6,250 cells/cm^2^; 1% O_2_; FBS supplementation, which exceeded the experimental threshold for 9 of 13 donors tested (Table 2). When ranked based on iDS, 5 of the 7 top-performing CPPs all shared low CD as a common parameter, while 4 of the 5 bottom-performing CPPs shared high CD O_2_ levels and MC did not show similar patterns, as predicted by the multi-response model (Fig 4A). The top performing CPPs (at low CD) favoured either FBS or hPL medium supplementation, while the bottom performing CPPs (at high CD) were agnostic to medium supplementation.

**Figure 4:**
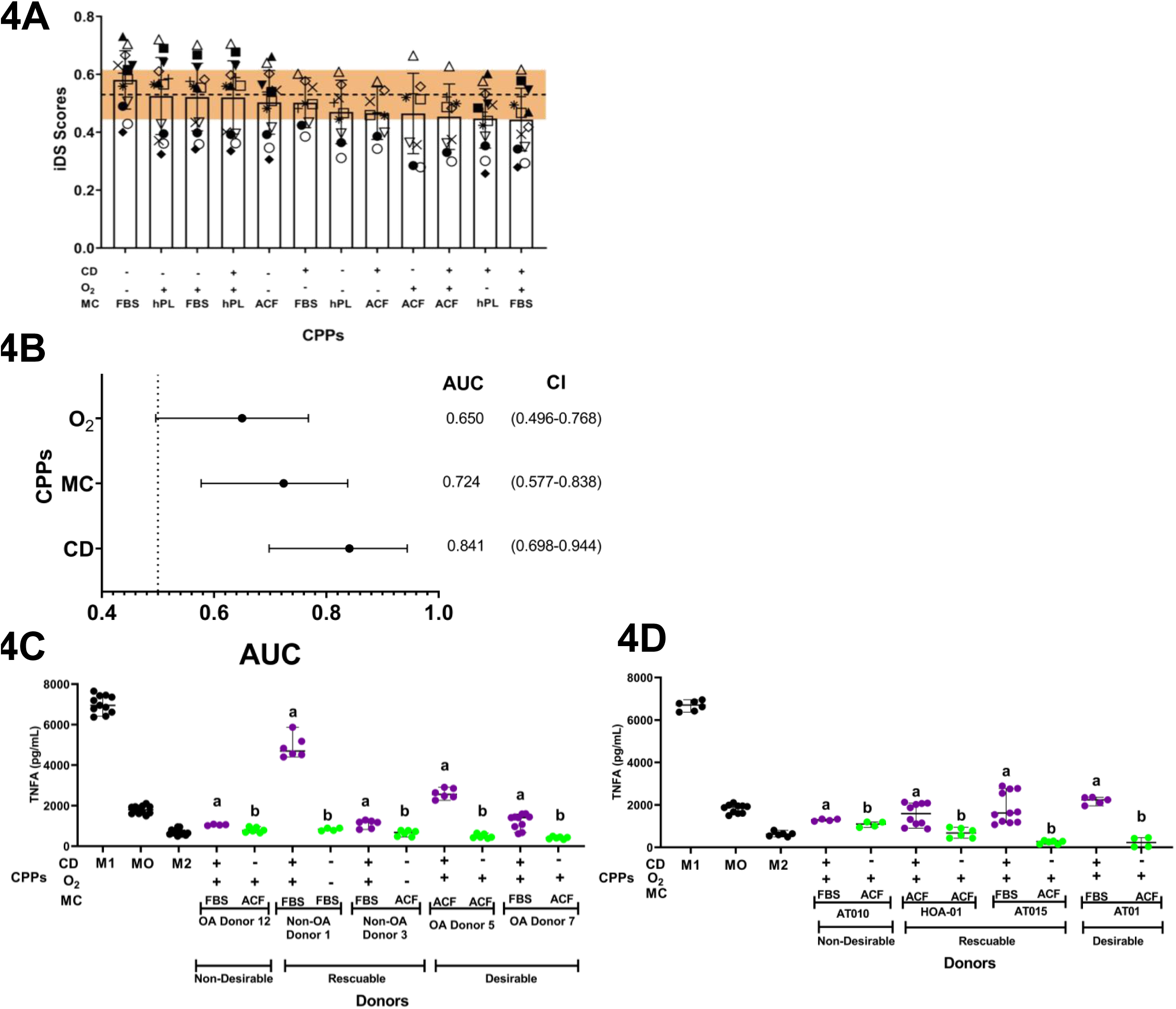
Model-predicted iDS across CPP conditions and independent functional assay validation for MSC(M) and MSC(AT) test and training datasets. (**A**) Model-predicted iDS using Model 1A across 12 CPP conditions and for all 13 MSC(M) training set donors. The experimentally determined iDS threshold (dotted line) at 0.530 ±0.086 (SD range shown in orange) is indicated. CPP combinations include CD, MC, and O_2_, with CD and O_2_ both evaluated at low (“–”) and high (“+”) levels; CD at 6,250 cells/cm² (“-”) and 31,250 cells/cm² (“+”); O_2_ at 1% (“-”) and 21% (“+”). (O) for Donor 1, (□) for Donor 2, (●) for Donor 4, (◼) for Donor5, (▲) for Donor 6, (□) for Donor 7, (□) for Donor 10, (▼) for Donor 11, (□) for Donor 12, (□) for Donor 13, (*) for Non-OA Donor 1, (+) for Non-OA Donor 2, and (X) for Non-OA Donor 3. (**B**) AUC analysis of O_2_, MC and CD for classifying TNF-reducing versus TNF-enhancing conditions depicted as Forest Plots with 95% CI values. Scatter plots of TNF (pg/mL) showing individual data points with mean and 95% CI from CD14□ monocyte-derived macrophages incubated with conditioned medium from (**C**) MSC(M) training or (**D**) MSC(AT) test dataset donors under CPP conditions associated with low predicted iDS (purple) or high model-predicted iDS (green). A minimum of 4 technical replicates were performed per condition, with additional replicates included where inter-replicate variability warranted. Controls (black) represent macrophages cultured in basal medium without MSC-conditioned medium (M0), or under pro-inflammatory (“M1”) or inflammation-resolving (“M2”) polarization conditions. Letters denote statistically significant differences between groups within each donor (p < 0.05). Letters indicate significant differences between specific groups using Krusal–Wallis non-parametric test.

Interestingly, only one donor (OA Donor 7) of 13 MSC(M) training set donors showed desirable model-predicted iDS across all 12 CPP combinations tested (Table 2). According to model predictions, a total of five donors showed desirable iDS across 9-10 of 12 CPP combinations tested; four of the bottom-performing donors were predicted to have undesirable iDS across all 12 CPP combinations tested; the remaining four intermediate donors (including all non-OA donors) showed CPP dependent iDS (Table 2). The top and intermediate performing donors favored CPP conditions associated with low seeding density, paired with hPL or serum-supplementation, while the bottom performing donors failed to exceed desirability thresholds, even at a low seeding densities.To test these model predictions, an independent functional assay measuring TNF secretion by primary CD14^+^ monocytes, differentiated to macrophages and exposed to MSC(M) conditioned medium (CM) was conducted. A subset of five MSC(M) donors (representing model predicted top, bottom and intermediate ranked donors from Table 1) were grown in different CPP conditions, reflecting high or low iDS, as predicted by the model (Suppl. Table 15). TNF levels were inversely correlated with model predicted iDS, associated with optimal or sub-optimal CPP conditions independently confirming model predictions. Specifically, CD, MC and O_2_ were evaluated for their ability to discriminate binary TNF outcomes, calculated against untreated or cytokine treated macrophage controls (Fig 4B). Of these three CPPs, CD demonstrated the strongest discrimination with an AUC of 0.841 (95% CI 0.70 – 0.94), with low CD conditions strongly associated with reduced macrophage TNF production. This aligned with iDS-based model predictions of top CPP conditions, which shared low CD as a common parameter. MC showed moderate but discrimination with an AUC of 0.724 (0.58 – 0.84), with ACF medium favouring reduced macrophage TNF relative to FBS/hPL-supplemented conditions; this slightly diverged from the model predictions, which did not favour any MC condition; similarly changes in O conditions resulted in low discriminatory power, consistent with model iDS ratings.

Model-predicted improvements in iDS translated into measurable differences in secondary, macrophage TNF production for individual MSC(M) training set donors (Fig. 4C). MSC(M) donors cultured under CPP combinations associated with higher iDS values resulted in lower TNF levels, concordant with cytokine polarized pro-resolving macrophage controls; model predicted, lower iDS associated CPP combinations for the same donors resulted in elevated TNF levels. Notably, MSC(M) training set donors ranked as high or intermediate potency performers showed greater CPP-dependent differences in TNF production by treated macrophages compared to donors ranked as lower performers. Thus, our model’s discriminative resolution was strongest when CPP optimization resulted in greater biological effects.

Further validation was conducted in an independent test dataset of five MSC(AT) donors, subjected to optimal and sub-optimal CPP combinations (Suppl. Table 15). As with the MSC(M) training dataset, model predictions of iDS, associated with a specific CPP combination were verified by inverse TNF production levels across all donors (Fig 4D). Additionally, as with the training dataset, there were greater changes in TNF production in donors that the model ranked as high or intermediate performers. Together, these data confirm the multi-response model predictions of optimal and sub-optimal CPP combinations in an independent functional assay.

## 4 Discussion

We developed a multi-response model to calculate a composite score iDS based dually on gene expression and doubling time as surrogate readouts of MSC quality attributes to evaluate the interplay between donor heterogeneity and CPPs. The multi-response model predicted iDS ranked 13 MSC(M) training set donors; donor rankings for a subset of 10 donors coincided with clinical responder categorization based on both significant Spearman’s correlations and high-performance AUC, verifying the utility of the model in predicting potent donors. Model predictions of potent donors were also tested in five MSC(AT), which coincided with previous potency rankings^28^.

The multi-response model was also used to assess the dichotomy of optimizing MSC expansion vs. surrogate potency attributes. This tension is well recognized in the field, yet few computational tools quantify where this trade-off limit lies, a challenge compounded by substantial donor heterogeneity inherent to the use of primary tissue derived MSCs. According to our model predictions, the best correlations to donor performance based on limited clinical data^27^ were generated by models that empirically weighted the cell expansion at 0%, 10% and 20% and correspondingly weighted gene expression at 100%, 90% or 80%. The top performing models thus disproportionately weighted MSC(M) gene expression over cell expansion. These findings may be specific to an autologous setting, dually capturing donor and host heterogeneity effects ^40^.

Even within these top performing models, 80% of model predicted top CPP combinations (across all donors) included low CD conditions, which resulted in high composite cell expansion and surrogate potency scores, consistent with reported effects of low seeding density on MSC proliferation^41^ and immunodulatory gene expression^42^. The model predicted that low seeding densities paired with serum-free or serum-supplemented conditions at all O_2_ concentrations was most effective in reliably producing high quality MSCs, up to 62-69% of MSC(M) training set donors. Conversely, the use of high seeding density, regardless of medium supplementation or O_2_ levels were the lowest ranked CPP conditions for 77% of MSC(M) training set donors tested. This seeding density effect was confirmed in a smaller MSC(AT) test dataset. Model predictions for both training and test datasets were further validated by an independent immunomodulatory assay that show reduced or increased TNF production by mismatched peripheral macrophages incubated with conditioned medium from MSC donors grown under low or high seeding density conditions respectively. These are striking findings; changing the seeding density of MSCs directly influenced the functional profile of secondary CD14^+^ monocyte effectors exposed to soluble factors from these MSCs.

While hypoxia is known to increase both MSC expansion^43^ and potency attributes^44^, model predictions, matched by secondary immune effector validation showed limited effects of changing O_2_ concentrations. We did note that MSC(M) training set immunomodulation genes showed dependence on O_2_ concentration, confirming previously reported roles^45,46^. Interestingly, the top performing CPP condition included both low MSC seeding density and hypoxic conditions, and was optimal for 9 of 13 donors tested, in agreement with previous data showing reduced senescence under combined low seeding density and hypoxia conditions^45^.

Based on predictions from the top performing model, only one donor (out of 13) was consistently predicted to exceed the pre-set iDS threshold across all CPPs. Approximately 62% of donors (8/13) were predicted to have high iDS only under certain, but not all CPP conditions tested, illustrating interplay between donor heterogeneity and CPP growth conditions. Across these donors, 34% of 96 donor-CPP combinations evaluated showed that low seeding density was the strongest driver. O_2_ tension had a modest effect with an equal number of donors favouring both normoxic vs. hypoxic conditions. Among medium supplementation, FBS and hPL were comparably favorable, more so than serum-free ACF; this may reflect a bias in the curated gene panel to be responsive to serum supplementation^47,48^. However, xenofree and chemically defined formulations have also been shown to preserve MSC immunosuppressive function relative to serum-supplemented contols^49^, aligned with our observations of better performance of serum-free conditions (over serum-supplementation) in modulating MSC impact on secondary macrophages. Divergence in model predictions for medium formulations, based on composite scores vs. secondary immunomodulatory functional assays, reflects the differential effects of a multitude of growth factors present on specific MSC potency attributes^50^. Overall, our model predicts that the majority of MSC(M) donors responded well in terms of both composite gene expression/cell expansion scores and functional immunomodulatory performance to low seeding density; however FBS or serum-free formulations, normoxia or hypoxia as optimal conditions varied from donor-to-donor.

Our model also predicted that approximately 30% of donors had low iDS and were predicted to perform poorly under all CPP conditions tested. Prospective identification and elimination of sub-performing donors that cannot be rescued even with CPP optimization thus becomes even more critical. Prospective donor identification based on gene expression has been proposed, based on inversely correlated *TWIST1 and TSG6* expression that identify high angiogenic vs. immunomodulatory potency donors^36,51–52^. This inverse relationship between angiogenic and immunomodulatory genes (specifically *PDCD1LG2, IDO and TNFAIP6*) was also demonstrated in our dataset with high performing donors showing increased immunomodulatory licensed gene expression at the expense of reduced angiogenic gene expression; immunomodulatory gene expression was also reduced as function of hypoxic conditions, aligned with this orthogonal relationship^36^.

Broadly, top-performing donors were predicted to be more robust, maintaining high composite gene expression/cell expansion across most CPP combinations, whereas low and intermediate performing donors achieved desirable outcomes for only a handful of CPP combinations. In fact, for the top five performing donors from the MSC(M) training dataset,75-100% of CPP combinations were predicted to have minimal impact on iDS. However, utilizing sub-optimal culture conditions was still predicted to reduce potency and expansion attributes even for top-performing donors, resulting in dramatic increases in TNF levels. Interestingly, the model predicted that lower potency donors were relatively insensitive to CPP conditions; their composite cell expansion and potency attributes could not be “rescued” to exceed the threshold even with changes to CPP conditions. This was confirmed in an independent functional assay which showed relatively minor, albeit significant fluctuations in TNF levels when CPP conditions were changed from optimal to sub-optimal. These results further stress the importance of screening donors and eliminating low performing donors that show minor improvements in potency attributes even with CPP optimization. Potent donors need to be continuously maintained in optimal conditions, and strong process controls at each stage should verify that potency attributes are not impacted by drifts in culture parameters^53^.Current in silico models for optimizing MSC and other cell-based therapies typically focus on maximizing cell expansion but often overlook potency optimization^54–55^. Donor variability is also often not integrated into these modeling approaches, but when integrated, provides valuable insights^56^. Artificial intelligence (AI)/Machine Learning (ML) models have the capacity to explicitly account for donor variability as recently shown^57^. However, most ML models require large, high-quality datasets that remain costly and labor intensive to generate. Here our multi-response approach provides a useful compromise, allowing us to sample over 1,000 multivariable readouts in approximately 100 experimental runs; we can capture the effects of the CPPs and their interactions with donor heterogeneity on dual MSC attributes, which cannot be transparently investigated in current ML models.

Advances such as Bayesian Optimization (BO) offer a more flexible alternative to the multi-response model approaches by employing surrogate models that can accommodate complex, non-linear effects and interactions without assuming a fixed response surface structure. BO iteratively updates its model of the design space and dynamically balances exploration of new conditions with exploitation of promising regions. This adaptability also allows the parameter space and sampling strategy to evolve across iterations, making BO particularly well-suited to the high-dimensional, donor-variable landscape of optimizing MSC attributes^58^.

Limitations to our model include the relatively modest sample size with thirteen MSC(M) training set donors and five MSC(AT) test dataset donors. Leave-out-one sensitivity analysis were applied to mitigate some aspects of the limited datasets. The parametric space was limited to three main parameters tested at multiple levels; additional parameters such as MSC adhesion, surface coating, medium change frequency, cryopreservation, and isolation methods were not considered but are known contributors to influencing MSC potency^31^. Using ML methods such as Gaussian process regression can expand this framework without the interaction-term constraints of multi-response surface models. However, these methods operate outside a hypothesis-testing framework and therefore do not directly address the uncertainty introduced by donor-to-donor variability; a model trained on limited donor data will carry large predictive uncertainty regardless of the optimization strategy employed and will require careful consideration.

Another limitation of the current model was the modest range of cell expansion (∼ 4-fold), which falls below clinical-scale MSC(M) manufacturing ranges (10-60 fold)^41,59^; thus, model predictions may not hold at larger scales and requires further verification. The smaller scale was chosen to allow relatively higher throughput experimental exploration. Importantly, model predictions of potent donors correlated with clinical efficacy; model-predicted optimal CPP combinations improved functional assay readouts in both training and test datasets, also increasing confidence in the scale used.

Together, our computational and experimental framework provide a meaningful tool for navigating MSC donor heterogeneity. Model-predicted potency and expansion scores correlated with clinical efficacy, and most donors (∼60%) demonstrated CPP-responsive immunomodulatory attributes, confirming that process optimization is a viable strategy for improving therapeutic consistency across much of the donor pool. Specifically, transitioning to low seeding density, and selecting serum vs. hPL conditions or O_2_ levels at the donor level results in optimized MSC composite quality attributes. Nonetheless, approximately 30% of tested donors exhibited persistently low potency scores, unresponsive to CPP adjustments, representing an intrinsic biological ceiling, especially in a diseased, autologous setting that process changes alone cannot overcome; this subset of donors may be prospectively identified through reduced expression of immunomodulatory genes. Collectively, these results support a path towards rational MSC manufacturing in which donor stratification and donor-dependent and predictive CPP optimization work in concert to improve potency considerations and can putatively reduce the risk of using sub-performing MSC batches.

## Supporting information

Graphical Abstract

## Acknowledgements

This research was funded by the Canadian Institutes of Health Research (CIHR), Natural Sciences and Engineering Research Council of Canada (NSERC, RGPIN 2018-05737), Data Science Institute (DSI DSFY1R1P12), This work was also funded by Toronto General and Western Hospital Foundation. We gratefully acknowledge the Schroeder Arthritis Biobank run by Dr. M Kapoor and assistance from K. Perry, C. Ward and L. Montoya. We thank M. Rasti and K. Fan for valuable assistance with the TNF assay.

COI

SV owns 60% of Regulatory Cell Therapy Consultants Inc; KR is currently employed by Stem Cell Technologies.

**Experimental and statistical framework to investigate interplay between critical processing parameters (CPPs) and donor heterogeneity on MSC(M) potency and cell expansion attributes**

<u>M</u>arrow MSC(<u>M</u>) (training dataset) were analyzed using a multi-response model to evaluate the interplay between donor heterogeneity and CPPs (CD, MC, and O□) on MSC(M) surrogate CQAs represented by gene expression and cell expansion. Various weightings were considered in converting gene expression and cell expansion experimentally-determined values into integrated desirability scores (iDS), across donors and CPP test conditions. Model predicted iDS were used to rank donors; donor ranking was independently validated against previous clinical effectiveness^27^; model predicted iDS was also used to rank CPP combinations; these were tested against an independent immunomodulatory functional assay. The test dataset comprised of five <u>a</u>dipose tissue MSC(AT) donors. ΔCT- delta cycle threshold; *ACF- Animal component free; CD- cell density; CPPs- critical process parameters; CQAs- critical quality attributes; ELISA-Enzyme-Linked Immunosorbent Assay; FBS- Fetal bovine serum; hPL- Human platelet lysate; iDS-Integrated desirability scores; KOOS- Knee Injury Osteoarthritis Outcome Score; MC-medium supplementation; MSC(AT)- Adipose Tissue MSC; MSC(M)-Bone Marrow MSC; OA-osteoarthritis; O_2_- oxygen concentration; RSM-response surface methodology;* T*_d_*-Doubling Time.

