## Supplementary figures and images for "Combined computational and experimental analysis confirm donor-dependent optimization of critical processing parameters for improving mesenchymal stromal cell potency and expansion attributes"

### Graphical Abstract

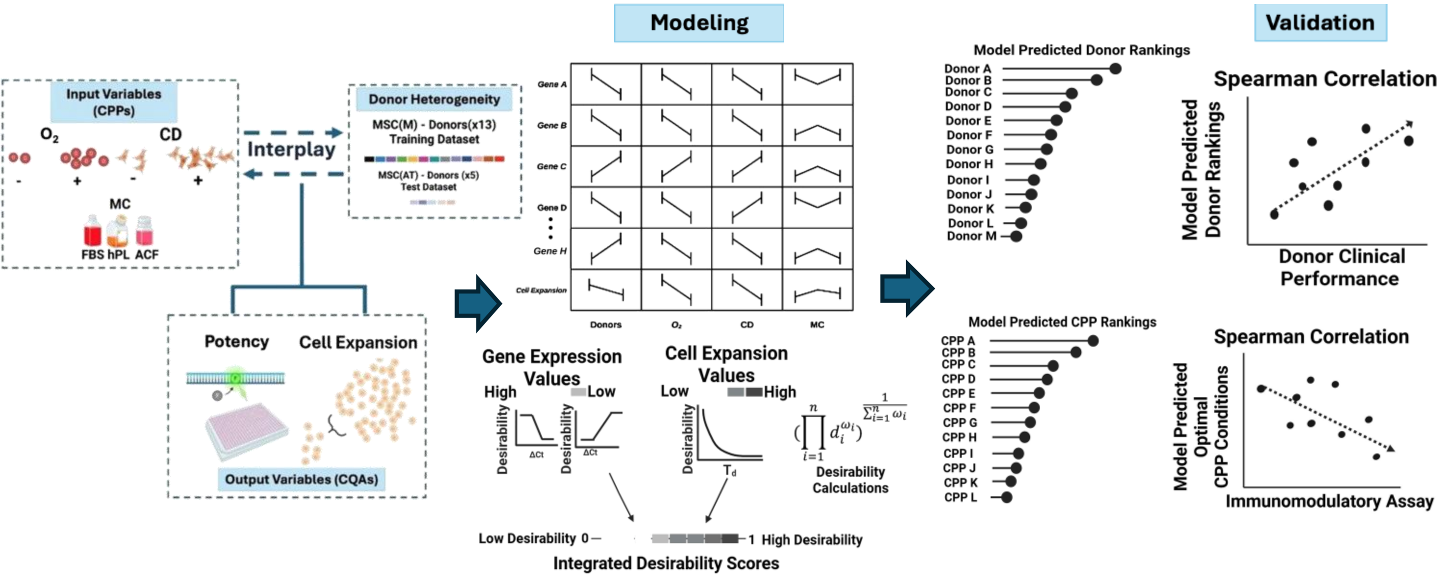
