## Supplementary files for "Combined computational and experimental analysis confirm donor-dependent optimization of critical processing parameters for improving mesenchymal stromal cell potency and expansion attributes"

**Supplementary Methods**

**MSC CPP Variables**

Donor MSC(M)s were cultured under three main variable critical process parameters (CPP) conditions: cell density (CD), medium composition (MC), and oxygen concentration (O_2_). Cells were seeded at either 6,250 cells/cm^2^ (low density) or 31,250 cells/cm^2^ (high density). The low density represents median cell density commonly used for MSC in monolayer expansions**^1^**, while the high density corresponds to the lower range of seeding densities utilized in MSC bioreactor systems**^2^**. MC was tested using three conditions:(i) 10% fetal bovine serum (FBS); (ii) human platelet lysate (hPL, STEMCELL Technologies) supplemented with heparin (Sterinova); or (iii) Mesencult ACF Plus supplemented with L-glutamine (Gibco) and Mesencult ACF Plus 500X supplement. O_2_ concentration was controlled using an Oxford Optronix HypoxyLab™ (1% hypoxic O_2_) or normoxic (21% O_2_) conditions using 5% CO_2_ ambient air according to previous protocols**^3^**.

**MSC(M) CQAs for Cell Expansion and Gene Expression**

MSC(M) and MSC(AT) were thawed and expanded as before**^3^** but substituted with DMEM low glucose (Sigma-Aldrich). Cell expansion was assessed after 7 days in different CPP conditions in 6-well plates, with medium changes performed every 3 days. Viable cells were recorded using a Vi-Cell Xr (Beckman Coulter) with confirmational counting by manual hemocytometer. Cell expansion, expressed as doubling time (T_d_), was calculated by dividing the expansion over 7 days by log base 2 of the cell expansion. As an outlier analysis, T_d_ was compared across Responders, Non-Responders, and non-OA donors over multiple ranges (0–10, 0–20, and 0–30 days) to assess whether outlier exclusion influenced the comparison. Within the 0–10 day range, Responders trended towards a shorter mean Td (3.79 ± 1.91 days) relative to Non-Responders (4.38 ± 2.16) and non-OA donors (4.16 ± 1.94), though no significant differences were observed between groups across any range tested. MSC(M)s were dually licensed with interferon-gamma (IFNy;100ng/mL) and tumor necrosis factor-alpha (TNF;10ng/mL) for 24h to enhance gene expression readouts as routinely conducted**^4^**. MSC(AT)s however were licensed using three cytokines,100ng/mL IFNg, 10ng/mL TNF and 5ng/mL interleukin (IL)-1 to benchmark against previously generated MSC(AT) gene expression data**^3^**. Total RNA was extracted using Ambion Trizol Reagent following the manufacturer’s protocol and quantified using a NanoDrop 1000 spectrophotometer. cDNA synthesis was carried out using VILO Master Mix (Invitrogen) according to manufacturer’s instructions. Gene expression was performed using Taq Pro Universal SYBR qPCR Master Mix (Vazyme) with specific primers for 8 MSC(M) curated gene expression (**Suppl. Table 2**). Raw cycle threshold (CT) values were calculated using the QuantStudio 5 system (Thermofisher). Gene expression was analyzed using the ΔCt method as previously described**^3^**, normalizing target gene expression against reference genes (*B2M, TBP*, and *GAPDH*). For PCA, negative ΔCt values (−ΔCt) were used as input, as higher values directly reflected higher expression and are more biologically intuitive. Raw ΔCt values were retained for downstream analysis and converted into individual desirability scores.

**MSC Donor Heterogeneity and CPP Interplay - Multi Response Models**

A first step in the multiple response analysis was to conduct factorial design experiments to systematically investigate the individual and interactive effects of donor-intrinsic parameters and CPPs during culture. This provided a training dataset of MSC(M), from different donors, cultured in a series of CPP combinations with corresponding ΔCt and T_d_ readouts; ΔCt values across eight curated genes and cell expansion measured as T_d_ together constituted potency attributes. The factorial design experiment was initially executed as a split-plot design with “Donor” treated as a random whole-plot factor to account for inter-donor variability. The design was subsequently augmented to incorporate later additions of new process parameters as an additional batch, and the combined dataset was analyzed with batch effects (Batch 1, Batch 2) included as an additional variable or blocking factor. Each response was used to fit a model composed of the same series of functional terms (**Equation 1**) which reflected a three-factor ANOVA with an additional blocking factor that accounted for the variation from the batch effect.s The model included all possible two-factor interactions between the CPPs.

*Y_h,i,j,k_ =β_o_ + S_h_ + β_i_ x_i_ + β_j_ x_j_ + β_k_ x_k_ +β_ij_ (x_i_ x_j_) + β_ik_ (x_i_x_k_) + β_jk_ (x_j_ x_k_) + e_h,i,j,k_* (**Equation** **1**)

Where Y_h,i,j,k_ corresponded, as applicable, to the transformed or non-transformed values of a given response (ΔCt values or T_d_) under specific combinations of donor and CPP conditions. β_o_ corresponded to the intercept; S_h_ was the blocking parameter for the batch effect (*h* = 2 categorical levels for two experimental levels ); *β_i_*, *β_j_,* and *β_k_* were the main effect coefficients for CD, MC, and O_2_ respectively; *β_ij_*, *β_i_*_k_, and *β_jk_* were the two-factor interaction coefficients; and x_i_, x_j_, x_k_ corresponded to the two categorical levels for each factor; ε corresponded to a normally distributed residual.

β coefficients in the model were estimated using the standard least square method and fitted for eight ΔCt values and T_d;_ statistical significance was assessed by F-test (p < 0.05). The F-test yielded a p-value for each coefficient, which was transformed into a log worth value (-log(pvalue)). When the homoscedasticity assumption was violated and/or the residuals could not be assumed to be normally distributed (based on the Shapiro-Wilk test), a Box-Cox transformation**^5^** was applied.

**Equation 2** and **3** applied the same framework as Equation 1 to Box-Cox transformed responses (*TGFB* +1, λ =0.5; *PDCD1LG1* +2, λ =0; *IDO1* +5, λ =0), and non-transformed responses (*VEGF, PDCD1LG2, TNFAIP6, TWIST1, HIF-1α,* T_d_) respectively, where g denoted each response variable fitted as an independent model.

*Y_g_=* *β_o_ + S_h_ + β _i_ x_i_ + β _j_ x_j_ + β _k_ x_k_ + β_ij_ (x_i_ x_j_) + β_ik_ (x_i_ x_k_) + β _jk (_x_j_ x_k_) + e_h,i,j,k_* (**Equation 2**)

Where g $\in$ {*TGF-B, PDCD1LG1, IDO1*}

*Y_g_* = *β_o_ + S_h_ + β _i_ x_i_ + β _j_ x_j_ + β _k_ x_k_ + β_ij_ (x_i_ x_j_) + β_ik_ (x_i_ x_k_) + β _jk (_x_j_ x_k_) + e_h,i,j,k_* (**Equation 3**)

Where g ∈ {*VEGF, PDCD1LG2, TNFAIP6, TWIST1, HIF-1a*, T_d_}

**Desirability Functions**

**Equation 1-3** were fitted by standard least squares method to simultaneously optimize CPP conditions across all nine MSC(M) potency responses. Individual desirability functions (d_r_, r = 1 to 9) fitted using the Derringer-Suich Desirability Function method**^6^** were developed for each response (**Equations 4-6**). ΔCt responses were adjusted for minimization (**Equation 4**) or maximization (**Equation 5**) to obtain DS_gene_ scores; while a ninth desirability function (f^9^, **Equation 6**) assigned T_d_-based desirability scores (DS_cell expansion_) to capture MSC(M) expansion across CPPs. For each desirability model, the maximum or minimum possible DS_cell expansion_ or DS_genes_ was numerically solved by computationally manipulating the levels of each of four factors (i.e. x_i_, x_j_, x_k_ and S_h_). This was performed using a gradient descent algorithm**^7-8^** to identify either the minimum (high expression of genes; low T_d_) or the maximum (low expression of genes; high T_d_) values for each CQA response across the training dataset. The desirability analysis was performed in JMP® (Pro 18) with the Profiler platform. DS_cell expansion_ and DS_genes_, derived from experimentally obtained T_d_ and ΔCT values and accounting for main and interaction effects, were integrated using weighted contributions for cell expansion and gene expression respectively into iDS values (**Equation 7, Suppl. Table 7**).

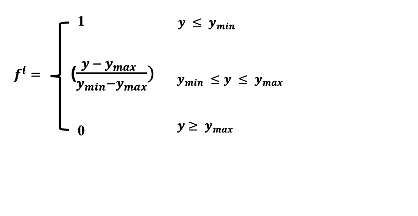
**(Equation 4)**

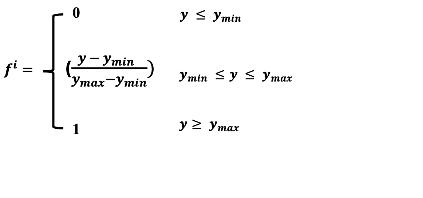
**(Equation 5)**

Where f^i^ denoted the individual desirability value for response y; y was the measured ΔCT response; y_min_ was the lowest experimentally measured ΔCT; and y_max_ was the highest experimentally measured ΔCT value, defined separately for each gene.

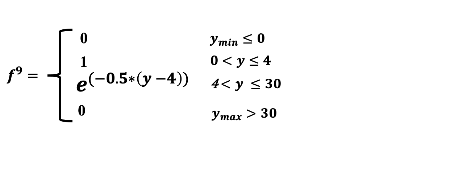
**(Equation 6)**

Where y_min_ and y_max_ represented the lowest and highest T_d_ values across all training set MSC(M) donors and CPPs. T_d_ ≤ 0 or > 30 days were assigned zero desirability; values between 0 and 4 days represented the optimal T_d_ range (f^9^ = 1) consistent with reported MSC doubling times**^9^**;T_d_ values from 4 to 30 days followed an exponential decay (e^-0.5(y-4)^) to capture the decreasing desirability of prolonged T_d_.

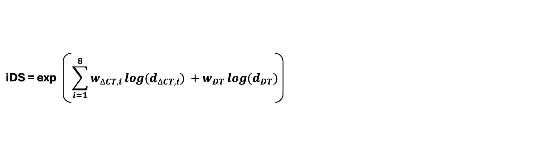
 **(Equation 7)**

Where *d_ΔCT,i_* denoted the desirability value for each gene (from **Equations 4 and 5**); *d_Td_* represented the T_d_-based desirability value (**Equation 6**); and *w_i_* and *w_Td_* are the per-gene and T_d_ weighting coefficients respectively.

**Composite desirability models incorporating multiple weighting strategies**

Thirteen models were generated by evaluating five ΔcT:T_d_ weighting schemes: 90:10 (Model 1), 80:20 (Model 2), 50:50 (Model 3), 100:0 (Model 4), and 0:100 (Model 5) with further sub-weightings. For Models 1 through 4, which retained a gene expression component, three sub-weighting strategies were applied across the eight curated genes: Model A incorporated all eight genes; Model B was restricted to the top three most sensitive genes, based on sensitivity analysis **(Suppl. Table 6**); and Model C applied directional weighting, penalizing culture conditions associated with increased *HIF1A, TWIST1*, or *VEGFA* expression while rewarding upregulation of the remaining genes.

The T_d_ weighting in each primary scheme reflected a distinct assumption regarding the relative importance of cell expansion. A 10% T_d_ weighting (Model 1) treated T_d_ as one of nine equally weighted variables alongside the eight genes. A 20% weighting (Model 2) represented an intermediate configuration, and a 50% weighting (Model 3) assigned equal importance to T_d_ relative to all eight genes combined. Models 4 and 5 served as limiting cases, applying 100% weighting exclusively to gene expression or T_d_, respectively. For Model A, all eight immunomodulatory and angiogenic genes were included, and **Equation 4** was applied as a minimization function to yield lower interaction-coefficient adjusted ΔCT values, corresponding to higher gene expression across all genes. DS_genes_ outputs were weighted by each gene's coefficient of variation (CoV) and directionality (**Suppl. Tables 6-7**). Model B was restricted to the top three most sensitive genes ( CoV > 10%; **Suppl. Table 6**) and applied the same minimization function, representing the empirically minimum number of gene outputs capable of reliably characterising MSC potency. Model C incorporated all eight genes but assigned each a functional role designation, either immunomodulatory (−1, minimization) or angiogenic (+1, maximization), based on the prevailing paradigm that these two functions represent relatively orthogonal potency axes in MSCs**^10^**.

For all thirteen models, gene-specific desirability values were incorporated into the iDS framework using **Equation 7**, where d_ΔcT,i_ denoted the desirability value for each individual gene derived from **Equations 4–5**, and T_d_ represented the T_d_ -based desirability value derived from **Equation 6**. The contribution of each term to the composite score was scaled by its respective weighting coefficient, w_i_ for individual genes and w_Td_ for T_d_ (**Suppl. Table 7**).

**ROC-AUC Analyses**Receiver Operating Characteristics (ROC) analysis was used to generate Area-Under-Cure (AUC). First, model-predicted iDS were used as a continuous predictor to classify all 13 donors as Responders or Non-responders based on KOOS outcome data. A second sent of AUCs was generated to assess the discriminative ability of three CPPs (O₂, MC, CD) against a secondary, independent MSC potency validation assay based on binary TNF-α levels (above or below a median threshold of 1,795.75 pg/mL based on control, untreated CD14⁺ macrophage-secreted TNF levels), with each CPP coded as a binary predictor (favourable = 1, unfavourable = 0). AUC with 95% confidence intervals were calculated using the DeLong method^11^.

**Supplementary Figure Legends**

**Supplementary Figure 1. PCA for subset of MSC(M) training set donors**

(A) PCA scores plot showing separation of 10/13 MSC(M) training set donors based on previous clinical responsiveness data**^12^**,Function-Pain Responders (blue) and Non-Responders (purple) along Component 1 (62.5%) and Component 2 (21.6%) under one CPP condition, 6250 cells/cm^2^ , 21% and FBS (B) PCA vector loadings plot identifying gene contributors to principal components (C) PC1 scores stratified by donor status; lowercase letters (a,b) denote statistically significant differences (p < 0.05, one-way ANOVA, Tukey's HSD). (D) Contribution of eight curated genes to PC1 and PC2 % variance. (E) Gene expression across 10 MSC(M) donors under same CPP condition, showing significant differences between Function-Pain Responders vs. Non-Responders. Pairwise comparison done with Tukey's HSD, significant differences labelled as varying letters. (F) AUC analysis of Function-Pain Responder (solid green) vs. Non-Responder (dashed purple) donors based on a composite biomarker signature (*TGFB,VEGF, PDCD1LG1, PDCD1LG2, IDO*). The model achieved perfect predictive performance (AUC = 1.00). PCA of all CPP conditions showed clustering by (G) donor and O_2_; solid circles represent hypoxia levels (1% O_2_), and hollow circles indicate normoxic levels (21% O_2_); by (H) donor and CD; solid circles represent high CD levels (31,250 cells/cm²), and hollow circles indicate low CD levels (6,250 cells/cm²) and by (I) donor and MC; solid circles for FBS, hollow circles for ACF; crosses for hPL. Function-Pain Responders in blue; Non-Responders in purple; and non-OA donors in green.

**Supplementary Figure 2. Doubling time (T_d_) of MSC(AT) under varying culture process parameters.**

Doubling time (T_d_) was assessed in a test dataset of MSC(AT) donors (n=5) expanded across CPP conditions: O_2_ at 1% (–) or21% (+), CD at 6,250 cells/cm² (–) or 31,250 cells/cm² (+), and MC comprising ACF, hPL, or FBS. Data is represented as mean T_d_ ± SD, individual points represent biological replicates. Lowercase letters indicate statistically significant pairwise differences (p < 0.05), determined by one-way ANOVA with Tukey’s HSD post-hoc test.

**Supplementary Figure 1**

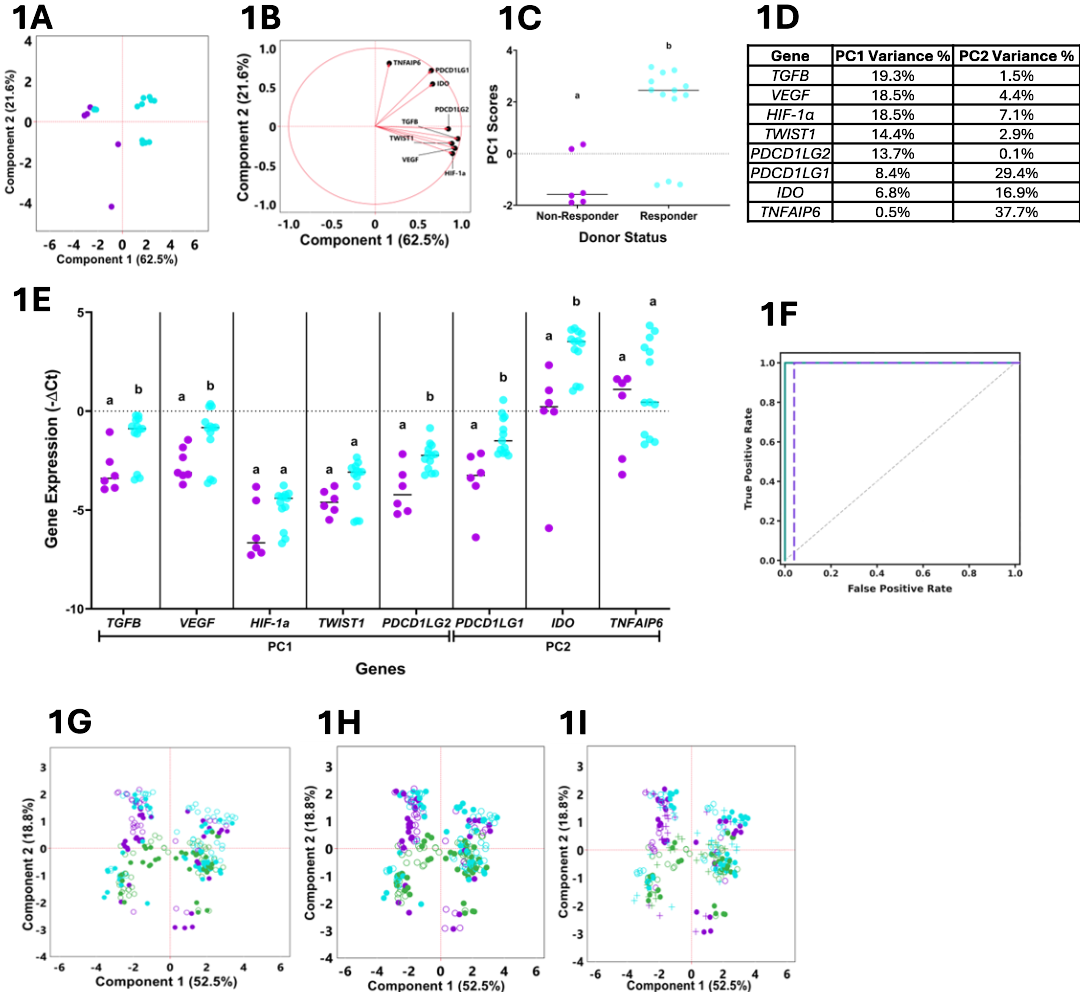

**Supplementary Figure 2**

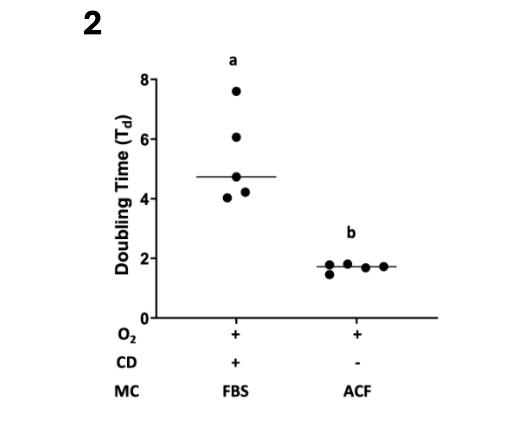

**Supplementary Tables**

**Supplementary Table 1:MSC(M) and MSC(AT) donor information.**

Donor characteristics including tissue source, age, sex, and BMI. N/A indicates data not available. *MSC(M)- bone marrow mesenchymal stromal cells; MSC(AT)- adipose tissue mesenchymal stromal cells; BMI- body mass index; OA – Osteoarthritis. *indicates missing donors, OA Donors 3 and 9 were not available in the biobank for analysis*

| **Dataset** | **Donor ID** | **Tissue** | **Age** | **Sex** | **BMI** |
| --- | --- | --- | --- | --- | --- |
| Training | **OA Donor***  **5,7,11,13** | MSC(M)  Function-Pain Responders | As reported in previously published data**^20^** | | |
| Training | **OA Donor*** **1,2,4,6,10,12** | MSC(M)  Function-Pain Non-Responders |  |  |  |
| Training | **Non-OA Donor 1** | MSC(M) | 32 | M | N/A |
| Training | **Non-OA Donor 2** | MSC(M) | 50 | M | N/A |
| Training | **Non-OA Donor 3** | MSC(M) | 58 | M | N/A |
| Test | **AT01** | MSC(AT) | 46 | M | 27.3 |
| Test | **AT07** | MSC(AT) | 28 | M | 26.9 |
| Test | **AT010** | MSC(AT) | 20 - 40 | F | N/A |
| Test | **AT015** | MSC(AT) | 52 | F | 22.4 |
| Test | **HOA-01** | MSC(AT) | 54 | M | 30.1 |

**Supplementary Table 2: Primers of MSC(M) and MSC(AT) curated gene panel**

*HIF-1a- hypoxia inducible factor 1 subunit alpha; IDO1- indoleamine 2,3-dioxygenase 1; PDCD1LG1- programmed cell death protein1 ligand 1; PDCD1LG2- programmed cell death protein 1 ligand2; TGFB1- transforming growth factor beta 1; TNFAIP6- tumor necrosis factor-inducible gene 6 protein; TWIST1- twist-related protein 1; VEGFA- vascular endothelial growth factor a*

| **Genes of Interest** | **Putative Categorization based on [Reference]** | **Forward Primers** | **Reverse Primers** |
| --- | --- | --- | --- |
| *TNF alpha-induced protein* 6  (*TNFAIP6*) | Anti-inflammatory/immunomodulatory^13^ | AGCACGGTCTGGCAAATACA | ATCCATCCAGCAGCACAGAC |
| *Indoleamine 2,3-dioxygenase* (*IDO1*) | Anti-inflammatory/immunomodulatory^14^ | GCCCTTCAAGTGTTTCACCAA | CCAGCCAGACAAATATATGCGA |
| *Programmed death-ligand 1*  (*PDCD1LG1*) | Anti-inflammatory/immunomodulatory^15^ | GCTGCACTAATTGTCTATTGGGA | AATTCGCTTGTAGTCGGCACC |
| *Programmed death-ligand 2*  (*PDCD1LG2*) | Anti-inflammatory/immunomodulatory^15^ | GTACATAATAGAGCATGGCAGCA | CCACCTTTTGCAAACTGGCTGT |
| *Transforming growth factor beta 1*  (*TGFB1*) | Anti-inflammatory/immunomodulatory^16^ | CTAATGGTGGAAACCCACAACG | TATCGCCAGGAATTGTTGCTG |
| *Twist-related protein 1*  (*TWIST1*) | Angiogenic^17^ | TCCATTTTCTCCTTCTCTGGAA | CCTTCTCGGTCTGGAGGAT |
| *Hypoxia-inducible factor 1-alpha*  (*HIF-1A*) | Angiogenic^18^ | ACGAGAGGTTCCCTAATTTCCA | ATGCCACCAGTACATTGGGAT |
| *Vascular Endothelial Growth Factor A*  (*VEGFA)* | Angiogenic^19^ | CTACCTCCACCATGCCAAGT | AGCTGCGCTGATAGACATCC |
| *Beta-2-microglobulin*  *(B2M*) | Housekeeping | CTTTCTGGCCTGGAGGCTATC | ACCAGTCCTTGCTGAAAGACAA |
| *TATA-binding protein*  (*TBP*) | Housekeeping | AGCGCAAGGGTTTCTGGTTT | AATAGGCTGTGGGGTCAGTC |
| *Glyceraldehyde-3-Phosphate Dehydrogenase*  (*GAPDH)* | Housekeeping | GGAGCGAGATCCCTCCAAAAT | GGCTGTTGTCATACTTCTCATGG |

**Supplementary Table 3: Doubling times (Td) of MSC(M) training dataset at different CPPs** MSC(M) training dataset doubling times (T_d_) in days under low CD (6250 cells/cm^2^) and high CD (31250 cells/cm^2^), cultured in hPL, ACF, or FBS supplemented medium at hypoxia (1% , "-") or normoxia (21%, "+") O_2_ . ND indicates runs excluded from the multi-response model which fell outside the planned DoE design space. Doubling time values >10 days were excluded as non-exponential and mathematically invalid, and correlation coefficients were agnostic to whether the exclusion threshold was set at 10 or 20 days. *ACF- animal component free; CD- cell density; CI-Confidence Interval; CPPs- critical processing parameters; IQR – Interquartile Range; O2- Oxygen; FBS- fetal bovine serum; hPL- human platelet lysate; MC- medium supplementation; Non-OA- non-osteoarthritis.*

|  | **Low CD** | | | | | | **High CD** | | | | | |
| --- | --- | --- | --- | --- | --- | --- | --- | --- | --- | --- | --- | --- |
|  | **hPL** | | **ACF** | | **FBS** | | **hPL** | | **ACF** | | **FBS** | |
| **Donor IDs** | - | + | - | + | - | + | - | + | - | + | - | + |
| OA Donor 1 | ND | 2.97 | 2.30 | ND | 4.65 | ND | 4.34 | ND | ND | ND | ND | 8.55 |
| OA Donor 2 | 2.82 | 3.14 | 1.57 | 3.37 | ND | 6.59 | ND | 9.79 | 4.08 | ND | ND | ND |
| OA Donor 4 | 1.98 | ND | 2.45 | ND | ND | 4.61 | ND | 4.65 | ND | ND | ND | ND |
| OA Donor 5 | 2.11 | 1.99 | 2.97 | 2.83 | 4.03 | ND | 5.37 | 4.12 | ND | 6.08 | ND | 4.38 |
| OA Donor 6 | 2.53 | 3.12 | ND | 2.37 | 4.84 | 6.34 | 8.41 | 4.71 | 6.00 | ND | ND | ND |
| OA Donor 7 | 1.79 | 2.10 | 1.93 | ND | 3.25 | 9.85 | ND | 5.16 | ND | 5.03 | ND | 4.38 |
| OA Donor 10 | 2.34 | ND | ND | 2.26 | ND | 5.27 | ND | 5.11 | 8.60 | ND | ND | ND |
| OA Donor 11 | ND | 2.04 | 2.16 | ND | ND | 4.61 | 5.28 | ND | ND | ND | ND | ND |
| OA Donor 12 | ND | 2.03 | 2.72 | ND | 5.33 | ND | 4.68 | ND | ND | ND | ND | ND |
| OA Donor 13 | ND | 2.05 | 2.16 | ND | ND | ND | 5.36 | ND | ND | ND | ND | ND |
| Non-OA Donor 1 | 2.29 | 2.05 | 2.65 | 3.48 | 4.27 | 5.47 | 4.27 | 3.84 | 9.86 | ND | ND | ND |
| Non-OA Donor 2 | 4.50 | ND | ND | 3.65 | ND | 7.00 | ND | 3.86 | ND | ND | ND | ND |
| Non-OA Donor 3 | 2.68 | 1.61 | 3.48 | 2.76 | 2.64 | 2.88 | 6.61 | 3.58 | ND | ND | 5.47 | 6.72 |

**Supplementary Table 4: DoE data availability 13 MSC donors x 12 CPP conditions**

Data availability across 13 MSC donors (10 OA, 3 non-OA) x 12 CPP conditions. Gene expression and doubling time data were collected (✓) or not (✕) for a given donor–condition combination. The 12 CPP conditions represent a full factorial of medium composition (ACF, FBS, hPL), cell density ("-"= 6,250cells/cm²; "+" = 31,250 cells/cm²), and oxygen tension ("-" = 1% or "+"= 21% O₂).

| **OA Donors** | **ACF** | | | | **FBS** | | | | **hPL** | | | |
| --- | --- | --- | --- | --- | --- | --- | --- | --- | --- | --- | --- | --- |
|  | **Low CD** | | **High CD** | | **Low CD** | | **High CD** | | **Low CD** | | **High CD** | |
|  | **-** | **+** | **-** | **+** | **-** | **+** | **-** | **+** | **-** | **+** | **-** | **+** |
| **1** | **✓** | ✕ | ✕ | **✓** | **✓** | ✕ | ✕ | **✓** | ✕ | **✓** | **✓** | ✕ |
| **2** | **✓** | **✓** | **✓** | **✓** | **✓** | **✓** | **✓** | **✓** | **✓** | **✓** | **✓** | **✓** |
| **4** | **✓** | ✕ | ✕ | **✓** | ✕ | **✓** | ✕ | ✕ | **✓** | ✕ | ✕ | **✓** |
| **5** | **✓** | **✓** | **✓** | **✓** | **✓** | **✓** | **✓** | **✓** | **✓** | **✓** | **✓** | **✓** |
| **6** | **✓** | **✓** | ✕ | **✓** | **✓** | **✓** | ✕ | ✕ | **✓** | **✓** | **✓** | **✓** |
| **7** | **✓** | **✓** | **✓** | ✕ | **✓** | **✓** | ✕ | **✓** | **✓** | **✓** | **✓** | **✓** |
| **10** | ✕ | **✓** | ✕ | **✓** | ✕ | **✓** | **✓** | ✕ | **✓** | ✕ | **✓** | ✕ |
| **11** | ✕ | ✕ | ✕ | **✓** | ✕ | **✓** | **✓** | ✕ | **✓** | ✕ | **✓** | ✕ |
| **12** | **✓** | ✕ | ✕ | **✓** | **✓** | ✕ | ✕ | **✓** | **✓** | ✕ | **✓** | ✕ |
| **13** | ✕ | **✓** | ✕ | **✓** | ✕ | **✓** | ✕ | ✕ | **✓** | ✕ | **✓** | ✕ |
| **Non-OA Donor 1** | **✓** | **✓** | **✓** | **✓** | **✓** | **✓** | **✓** | **✓** | **✓** | **✓** | **✓** | **✓** |
| **Non-OA Donor 2** | ✕ | **✓** | ✕ | **✓** | ✕ | **✓** | ✕ | ✕ | **✓** | ✕ | **✓** | ✕ |
| **Non-OA Donor 3** | **✓** | **✓** | **✓** | **✓** | **✓** | **✓** | **✓** | **✓** | **✓** | **✓** | **✓** | **✓** |

**Supplementary Table 5: Genes and corresponding residual p-values used for normalization**

p-values of the Shapiro-Wilk test for the normal distribution of the residuals of the models to predict gene expression obtained following Box-Cox transformation. *DoE- Design of Experiments*

| **Genes** | **p-Value** |
| --- | --- |
| *TWIST1* | 0.8920 |
| *IDO* | 0.4828 |
| *VEGFA* | 0.1896 |
| *TGFB* | 0.0828 |
| *PDCD1LG2* | 0.0424 |
| *HIF-1a* | 0.0264 |
| *TSG-6* | 0.0084 |
| *PDCD1LG1* | 0.0008 |

**Supplementary Table 6: Sensitivity analysis across eight genes**

CoV measured for eight genes across 13 MSC(M) training dataset donors including OA and non-OA donors. Bolded values indicate CoV greater than 10%. *CoV- Coefficient of Variation*

| ***Genes*** | ***All OA Donors*** | ***OA Function-Pain Responder Donors*** | ***OA Non-Responder Donors*** | ***Non-OA Donors*** |
| --- | --- | --- | --- | --- |
| *TNFAIP6* | ***24.30*** | ***32.33*** | ***22.12*** | ***14.95*** |
| *VEGFA* | ***13.27*** | ***10.88*** | ***12.35*** | ***12.73*** |
| *PDCD1LG2* | ***10.09*** | *8.54* | ***10.74*** | *7.67* |
| *TWIST1* | *8.20* | *7.87* | *7.33* | *7.9 0* |
| *HIF1A* | *7.24* | *5.13* | *8.6 0* | *4.86* |
| *IDO1* | *6.93* | *4.3* | *5.64* | *7.76* |
| *PDCD1LG1* | *4.84* | *4.44* | *4.27* | *3.38* |
| *TGFB1* | *2.84* | *2.42* | *2.62* | *2.43* |

**Supplementary Table 7: Gene Expression (ΔCT) and Doubling times (T_d_) weights and directionality across thirteen iDS model simulations**

Weighting coefficients for ΔCT (w_i_) derived from sensitivity analysis (*Suppl. Table 6*) and T_d_ (wTd ), applied in Equation 7 (*Supplementary Files*). Weights were assigned based on gene directionality, CoV, and proportional distribution across five ΔCT-to-Td ratios, evaluated separately for immunomodulatory and angiogenic gene configurations. "+" and "−" denote a desired decrease or increase in ΔCT, respectively; since lower ΔCT reflects higher gene expression, "−" corresponds to a target of increased expression. *NA - Not applicable shows no contributions in the weighting scheme;* ΔCT*- delta cycle threshold; T_d_ -Doubling Time*

| **Genes** | **Model 1A** | **Model 1B** | **Model 1C** | **Model 2A** | **Model 2B** | **Model 2C** | **Model 3A** | **Model 3B** | **Model 3C** | **Model 4A** | **Model 4B** | **Model 4C** | **Model 5** |
| --- | --- | --- | --- | --- | --- | --- | --- | --- | --- | --- | --- | --- | --- |
| *TNFAIP6* | -0.28 | -0.46 | -0.28 | -0.25 | -0.41 | -0.25 | -0.16 | -0.24 | -0.16 | -0.31 | -0.51 | -0.31 | NA |
| *VEGFA* | -0.15 | -0.25 | 0.15 | -0.14 | -0.22 | 0.14 | -0.08 | -0.13 | 0.08 | -0.17 | -0.28 | 0.17 | NA |
| *PDCD1LG2* | -0.12 | -0.19 | -0.12 | -0.10 | -0.17 | -0.10 | -0.06 | -0.13 | -0.06 | -0.13 | -0.21 | -0.13 | NA |
| *TWIST1* | -0.09 | NA | 0.09 | -0.08 | NA | 0.08 | -0.05 | NA | 0.05 | -0.11 | NA | 0.11 | NA |
| *HIF1A* | -0.08 | NA | 0.08 | -0.07 | NA | 0.07 | -0.05 | NA | 0.05 | -0.09 | NA | 0.09 | NA |
| *IDO1* | -0.08 | NA | -0.08 | -0.07 | NA | -0.07 | -0.04 | NA | -0.04 | -0.09 | NA | -0.09 | NA |
| *PDCD1LG1* | -0.06 | NA | -0.06 | -0.05 | NA | -0.05 | -0.03 | NA | -0.03 | -0.06 | NA | -0.06 | NA |
| *TGFB* | -0.03 | NA | -0.03 | -0.03 | NA | -0.03 | -0.02 | NA | -0.02 | -0.04 | NA | -0.04 | NA |
| Doubling Time | -0.10 | -0.10 | -0.10 | -0.20 | -0.20 | -0.20 | -0.50 | -0.50 | -0.50 | NA | NA | NA | 1 |

**Supplementary Table 8: ΔCT values at 50% median for select genes and associated error** rate with classification of Function-Pain Responders vs. Non-Responders. ΔCT values for individual OA donor samples across four genes from a previously published dataset**^20^**. Error was calculated as the proportion of cases misclassified by a 50% median, where complete separation of Function-Pain Responders from Non-Responders yielded 0% error.

| **Donor** | ***TNFAIP6* ΔCT** | ***PDCD1LG1* ΔCT** | ***TGFβ* ΔCT** | ***IDO1* ΔCT** |
| --- | --- | --- | --- | --- |
| OA Donor 1 | 16.55 | 5.02 | 5.7 | 2.55 |
| OA Donor 2 | 15.91 | 3.66 | 3.66 | 0.66 |
| OA Donor 4 | NA | 3.35 | 5.15 | 1.25 |
| OA Donor 5 | 12.71 | -0.33 | 1.4 | -0.79 |
| OA Donor 6 | NA | NA | NA | 9.55 |
| OA Donor 7 | NA | 4.03 | 4.03 | 2.03 |
| OA Donor 10 | NA | 3.09 | 3.09 | 2.09 |
| OA Donor 11 | NA | 2.35 | 4.1 | 3.1 |
| OA Donor 12 | NA | 3.66 | 3.81 | 1.65 |
| OA Donor 13 | NA | -0.72 | 0.28 | -1.57 |
| Average 50% Threshold (ΔCT) | 14 | 2 | 3 | 2 |
| Error Rate | 0% | 22.2% | 22.2% | 50% |

**Supplementary Table 9: Regression equations converting raw ΔCT measurements to gene expression desirability scores for experimental threshold determination (DS_Gene Threshold_)**

| **Genes** | **50% median**  **ΔCT value** | **Box-Cox Transformation ΔCT** | **Regression Equations** | **DS_Gene Threshold_** |
| --- | --- | --- | --- | --- |
| *TNFAIP6* | -1. | -1. | – 0.0769(ΔCT*TNFAIP6* ) + 0.559 | 0.63 |
| *VEGF* *A* | 1.7 | 1.7 | -0.148(ΔCT*VEGF*) + 0.667 | 0.416 |
| *PDCD1LG2* | 2.8 | 2.8 | – 0.139(ΔCT*PDCD1LG2*) + 1.016 | 0.620 |
| *TWIST1* | 3.8 | 3.8 | – 0.158 (ΔCT*TWIST1*) + 1.158 | 0.549 |
| *HIF-1α* | 5.04 | 5.04 | – 0.186(ΔCT*HIF-1a*) + 1.577 | 0.630 |
| *Transformed IDO1*  (λ=0, Constant =5) | -5.121 | 0.886 | – 0.29(ΔCT*IDO1*) + 0.914 | 0.659 |
| *Transformed PDCD1LG1*  (λ=0, Constant =2) | -2 | 1 | – 0.301 (ΔCT*PDCD1LG1*) + 0.803 | 0.500 |
| *Transformed TGFB1*  (λ=0.5, Constant =1) | 1.56 | 1.6 | – 0.590 (ΔCT*TGFB1*) + 1.395 | 0.445 |
| Average | N/A | N/A | N/A | 0.556 |

**Supplementary Table 10: Doubling time (T_d_) conversion to desirability scores for cell expansion to calculate an experimental threshold (DS_cell expansion threshold_).** T_d_ (days) for a subset of MSC(M) training set donors 10/13 at passage 3 from previous dataset**^20^** converted to DS_cell expansion threshold_ using the regression equation *f(DT)* = 0.42 +7.24•10^-4^*DT*. *DS- Desirability Score; T_d_- Doubling Time*

| **OA Donors ID** | **T_d_ (Days)** | **DS_Cell Expansion_ _Threshold_** |
| --- | --- | --- |
| 1 | 4.84 | 0.429 |
| 2 | 2.77 | 0.427 |
| 4 | 2.50 | 0.427 |
| 5 | 3.29 | 0.427 |
| 6 | 2.54 | 0.427 |
| 7 | 2.44 | 0.427 |
| 10 | 1.70 | 0.426 |
| 11 | 2.31 | 0.427 |
| 12 | 2.31 | 0.427 |
| 13 | 4.77 | 0.428 |
| Average | 3.41 | 0.427 |

**Supplementary Table 11. Average iDS for training MSC(M) dataset donors using different modelling weighting strategies**

Model predicted average iDS with SD (sub-scripted) for MSC(M) training dataset donors across 13 models as detailed in S*upplementary Methods*. SD (0.086) were calculated for each donor by averaging across all CPPs (n =12) per model. Average iDS exceeding the threshold of 0.530 were bolded; values above the threshold (0.530) were classified as desirable, those below threshold - SD (0.444) as non-desirable, and those falling within the range of the experimentally determined iDS threshold - SD (0.444-0.530) as rescuable.

| **OA Donor ID** | **Model 1A** | **Model 2A** | **Model 3A** | **Model 4A** | **Model 1B** | **Model 2B** | **Model 3B** | **Model 4B** | **Model 1C** | **Model 2C** | **Model 3C** | **Model 4C** | **Model 5** |
| --- | --- | --- | --- | --- | --- | --- | --- | --- | --- | --- | --- | --- | --- |
| 7 | **0.65 _0.05_** | **0.62 _0.09_** | **0.58 _0.19_** | **0.67_0.05_** | **0.62 _0.05_** | **0.6 0_0.08_** | **0.56 _0.19_** | **0.61_0.07_** | **0.55 _0.07_** | **0.54 _0.10_** | **0.53 _0.19_** | **0.54_0.09_** | **0.56_0.33_** |
| 5 | **0.59 _0.07_** | **0.57 _0.09_** | **0.55 _0.18_** | **0.60_0.08_** | **0.56 _0.08_** | **0.55 _0.09_** | **0.54 _0.18_** | **0.56_0.09_** | **0.57 _0.07_** | **0.56 _0.09_** | **0.55 _0.19_** | **0.56_0.08_** | **0.57_0.33_** |
| 6 | **0.58 _0.08_** | **0.57 _0.10_** | **0.54 _0.19_** | **0.61_0.08_** | **0.57 _0.09_** | **0.56 _0.10_** | **0.53 _0.19_** | **0.56_0.10_** | **0.58 _0.06_** | **0.56 _0.09_** | **0.54 _0.19_** | **0.57_0.08_** | **0.54_0.33_** |
| 11 | **0.57 _0.05_** | **0.56 _0.08_** | **0.54 _0.18_** | **0.59_0.05_** | **0.53 _0.06_** | 0.52 _0.08_ | 0.52 _0.18_ | **0.53_0.07_** | 0.5 0_0.08_ | 0.5 0_0.10_ | 0.51 _0.18_ | 0.50_0.09_ | **0.56_0.33_** |
| 13 | **0.56 _0.06_** | **0.53 _0.11_** | 0.48 _0.21_ | **0.60_0.04_** | 0.5 0_0.06_ | 0.48 _0.10_ | 0.45 _0.21_ | 0.49_0.09_ | 0.42 _0.06_ | 0.41 _0.10_ | 0.41 _0.19_ | 0.41_0.08_ | 0.47_0.33_ |
| Non-OA Donor 2 | **0.53 _0.05_** | 0.47 _0.08_ | 0.5 0_0.19_ | **0.55_0.05_** | 0.47 _0.04_ | 0.47 _0.07_ | 0.47 _0.18_ | 0.47_0.06_ | 0.46 _0.05_ | 0.46 _0.08_ | 0.46 _0.18_ | 0.46_0.07_ | 0.52_0.33_ |
| 2 | 0.51 _0.05_ | 0.50 _0.08_ | 0.49 _0.18_ | 0.52_0.05_ | 0.46 _0.05_ | 0.45 _0.07_ | 0.46 _0.17_ | 0.45_0.06_ | 0.43 _0.06_ | 0.43 _0.08_ | 0.45 _0.17_ | 0.43_0.08_ | 0.52_0.33_ |
| Non-OA Donor 1 | 0.5 0_0.05_ | 0.51 _0.06_ | 0.52 _0.15_ | 0.51_0.06_ | 0.44 _0.05_ | 0.45 _0.06_ | 0.49 _0.15_ | 0.45_0.05_ | 0.45 _0.04_ | 0.45 _0.06_ | 0.49 _0.15_ | 0.45_0.05_ | **0.60_0.33_** |
| Non-OA Donor 3 | 0.47 _0.09_ | 0.52 _0.08_ | 0.52 _0.14_ | 0.46_0.10_ | 0.48 _0.08_ | 0.49 _0.08_ | **0.53 _0.15_** | 0.48_0.08_ | 0.47 _0.03_ | 0.48 _0.06_ | 0.52 _0.14_ | 0.47_0.05_ | **0.64_0.32_** |
| 10 | 0.41 _0.04_ | 0.41 _0.06_ | 0.44 _0.14_ | 0.40_0.05_ | 0.41 _0.05_ | 0.41_0.06_ | 0.45 _0.15_ | 0.41_0.05_ | **0.54 _0.05_** | **0.54 _0.08_** | **0.53 _0.18_** | **0.54_0.07_** | **0.55_0.33_** |
| 4 | 0.38 _0.05_ | 0.39 _0.06_ | 0.43 _0.13_ | 0.37_0.06_ | 0.39 _0.06_ | 0.4 0_0.07_ | 0.44 _0.15_ | 0.40_0.06_ | **0.6 0_0.04_** | **0.59 _0.08_** | **0.56 _0.19_** | **0.59_0.07_** | **0.56_0.33_** |
| 1 | 0.34 _0.04_ | 0.35 _0.06_ | 0.41 _0.13_ | 0.33_0.05_ | 0.29_0.05_ | 0.31 _0.06_ | 0.38 _0.13_ | 0.30_0.05_ | 0.45 _0.04_ | 0.45 _0.07_ | 0.47 _0.16_ | 0.45_0.06_ | **0.57_0.33_** |
| 12 | 0.31 _0.04_ | 0.32_0.05_ | 0.37 _0.13_ | 0.30_0.05_ | 0.3 0_0.05_ | 0.31 _0.06_ | 0.37 _0.14_ | 0.31_0.05_ | 0.42 _0.04_ | 0.42 _0.07_ | 0.44 _0.16_ | 0.42_0.06_ | **0.53_0.33_** |

**Supplementary Table 12. AUC performance summary for 13 predictive models, ranked from highest to lowest discriminative performance.**

Area under the receiver operating characteristic curve with the 95% confidence interval calculated using the DeLong method. Models are ranked by AUC from highest to lowest discriminative performance.

| **Model** | **Line Color/Line Style** | **AUC** | **95% CI** |
| --- | --- | --- | --- |
| Model 4A | Orange/Solid | 0.902 | 0.846–0.957 |
| Model 1A | Red/Solid | 0.891 | 0.833–0.949 |
| Model 1B | Red/Dotted | 0.870 | 0.804–0.935 |
| Model 2A | Blue/Solid | 0.849 | 0.777–0.920 |
| Model 4B | Orange/Dotted | 0.844 | 0.772–0.916 |
| Model 2B | Blue/Dotted | 0.825 | 0.749–0.901 |
| Model 3A | Green/Solid | 0.661 | 0.556–0.765 |
| Model 3B | Green/Dotted | 0.644 | 0.539–0.750 |
| Model 4C | Orange/Dashed | 0.521 | 0.414–0.629 |
| Model 2C | Blue/Dashed | 0.518 | 0.411–0.625 |
| Model 1C | Red/Dashed | 0.517 | 0.410–0.623 |
| Model 3C | Green/Dashed | 0.503 | 0.396–0.611 |
| Model 5 | Purple/Solid | 0.499 | 0.392–0.605 |

**Supplementary Table 13: Leave-out one analysis**

Spearman correlation coefficients and corresponding p-values were calculated between iDS (for top-performing Model 1A) and combined KOOS pain and function percent improvements at 12-months, relative to baseline**^12^** after omitting one donor (from a subset of 10/13 MSC(M) training set donors for which clinical performance data was previously available**^20^**) at a time as indicated in the top row. Statistically significant average correlations (*p<0.10) using Spearman’s correlation are bolded.

| Follow-up Point | KOOS  Scores | OA-Donor 1  Excluded | | OA-Donor 2  Excluded | | OA-Donor 4  Excluded | | OA-Donor 5 Excluded | | OA-Donor 6 Excluded | | OA-Donor 7  Excluded | | OA-Donor 10 Excluded | | OA-Donor 11 Excluded | | OA-Donor 12 Excluded | | OA-Donor 13  Excluded | | Leave -One-Out Average | | No exclusions | |
| --- | --- | --- | --- | --- | --- | --- | --- | --- | --- | --- | --- | --- | --- | --- | --- | --- | --- | --- | --- | --- | --- | --- | --- | --- | --- |
| 12  Month |  | Spearman  p | Prob > \|p\| | Spearman  p | Prob > \|p\| | Spearman  p | Prob > \|p\| | Spearman p | Prob > \|p\| | Spearman p | Prob  > \|p\| | Spearman  p | Prob  >\|p\| | Spearman p | Prob  >\|p\| | Spearman p | Prob  > \|p\| | Spearman  p | Prob  > \|p\| | Spearman  p | Prob  >\|p\| | Spearman  p | Prob  >\|p\| | Spearman  p | Prob  >\|p\| |
|  | Combined Percent | 0.67 | 0.05 | 0.77 | 0.02 | 0.57 | 0.11 | 0.67 | 0.05 | 0.87 | 0.003 | 0.62 | 0.08 | 0.73 | 0.02 | 0.47 | 0.21 | 0.60 | 0.09 | 0.65 | 0.06 | 0.66 | 0.07 | 0.72 | 0.02 |
|  | %Change ADL | 0.66 | 0.05 | 0.80 | 0.01 | 0.59 | 0.09 | 0.62 | 0.08 | 0.86 | 0.003 | 0.62 | 0.08 | 0.74 | 0.02 | 0.46 | 0.21 | 0.59 | 0.09 | 0.63 | 0.07 | 0.66 | 0.07 | 0.72 | 0.02 |
|  | %Change Pain | 0.87 | 0.003 | 0.85 | 0.004 | 0.73 | 0.02 | 0.88 | 0.002 | 0.83 | 0.01 | 0.78 | 0.01 | 0.85 | 0.004 | 0.73 | 0.02 | 0.77 | 0.02 | 0.83 | 0.01 | 0.81 | 0.01 | 0.84 | 0.002 |

**Supplementary Table 14: Gene Combination Rankings of MSC(AT) compared to previous adipose rankings.**

MSC(AT) rankings based on model predicted iDS across 3 conditions, compared to previous rankings.

| **Previous Donor Rankings^3^** | **Predicted Donor Rankings at 1%/6250/FBS** | | **Predicted Donor Rankings at 21%/6250/ACF** | | **Predicted Donor Rankings at 21%/31250/FBS** | |
| --- | --- | --- | --- | --- | --- | --- |
| **Donor** | **Donor** | **iDS** | **Donor** | **iDS** | **Donor** | **iDS** |
| AT01 | AT01 | 0.72 | AT01 | 0.55 | AT01 | 0.55 |
| AT015 | AT015 | 0.64 | AT015 | 0.51 | AT015 | 0.47 |
| AT07 | AT010 | 0.42 | HOA-01 | 0.43 | HOA-01 | 0.44 |
|  | HOA-01 | 0.37 | AT-010 | 0.41 | AT-010 | 0.41 |
|  | AT07 | 0.37 | AT07 | 0.32 | AT07 | 0.24 |

**Supplementary Table 15: Selected CPP combinations for MSC(M) training and MSC(AT) test donors based on top performing model (Model 1A) predicted iDS.**

MSC(M) training dataset donors were stratified into three categories based on their mean iDS: i) iDS that exceeded the threshold (0.530) = Desirable (bolded); ii) iDS with SD range [0.444, 0.530] ="Rescuable"; iii) iDS below 0.444= "Non-Desirable" For each donor category, optimal and suboptimal CPP combinations were identified that were associated with corresponding iDS (*from Table 2*).

| **Dataset** | **Ranking** | **MSC(M) Donors** | **High iDS** | | **Low iDS** | |
| --- | --- | --- | --- | --- | --- | --- |
|  |  |  | **iDS** | **Corresponding CPP** | **iDS** | **Corresponding CPP** |
| Training | Desirable | OA Donor 7 | 0.66 | 21,6250,ACF | 0.62 | 21,31250,FBS |
|  | Desirable | OA Donor 5 | 0.62 | 21,6250,ACF | 0.58 | 21,31250,FBS |
|  | Rescuable | Non-OA Donor 1 | 0.56 | 1,6250,FBS | 0.49 | 21,31250,FBS |
|  | Rescuable | Non-OA Donor 3 | 0.54 | 1,6250,ACF | 0.39 | 21,31250,FBS |
|  | Non-Desirable | OA Donor 12 | 0.34 | 21,6250,FBS | 0.28 | 21,31250,FBS |
| Test | **Ranking** | **MSC(AT) Donors** | **iDS** | **Corresponding CPP** | **iDS** | **Corresponding CPP** |
|  | Desirable | AT01 | 0.55 | 21,6250,ACF | 0.55 | 21,31250,FBS |
|  | Rescuable | AT015 | 0.51 | 21,6250,ACF | 0.47 | 21,31250,FBS |
|  | Rescuable | HOA-01 | 0.43 | 21,6250,ACF | 0.44 | 21,31250,ACF |
|  | Non-Desirable | AT010 | 0.41 | 21,6250,ACF | 0.41 | 21,31250,FBS |
|  | Non-Desirable | AT07 | 0.32 | 21,6250,ACF | 0.24 | 21,31250,FBS |

**Supplementary Files References**
